# GCL pruning of PIP3 establishes the soma-germline boundary

**DOI:** 10.64898/2025.12.30.697122

**Authors:** Mariyah Saiduddin, Juhee Pae, Asier M. Vidal, Marty Alani, Ruth Lehmann

## Abstract

Primordial germ cells (PGCs) are the first cells specified in the Drosophila embryo and serve as precursors to the germline. Their formation requires suppression of somatic fates, a process achieved by excluding the receptor tyrosine kinase Torso from the posterior pole through degradation mediated by the ubiquitin ligase adaptor Germ Cell-Less (GCL). Although Torso is known to antagonize PGC formation, the underlying mechanism has remained unclear. Here, we combine optogenetic Ras activation and Ras effector loop mutants to show that Ras signaling suppresses PGC formation independently of the canonical Raf/MEK/ERK pathway. We identify an unexpected early role for Torso in activating phosphoinositide 3-kinase (PI3K), generating posterior membrane domains enriched in phosphatidylinositol (3,4,5)-trisphosphate (PIP3). Elevated PI3K activity disrupts PGC formation, while reduced PI3K activity leads to ectopic PGCs. We further demonstrate that GCL remodels the posterior pole membrane by suppressing Torso-dependent PI3K activation. Clearing PIP3 enables Myosin II enrichment, thereby constricting the pole bud for PGC formation. Together, our findings reveal how antagonistic Torso and GCL activities establish the soma–germline boundary by regulating cortical lipid organization.

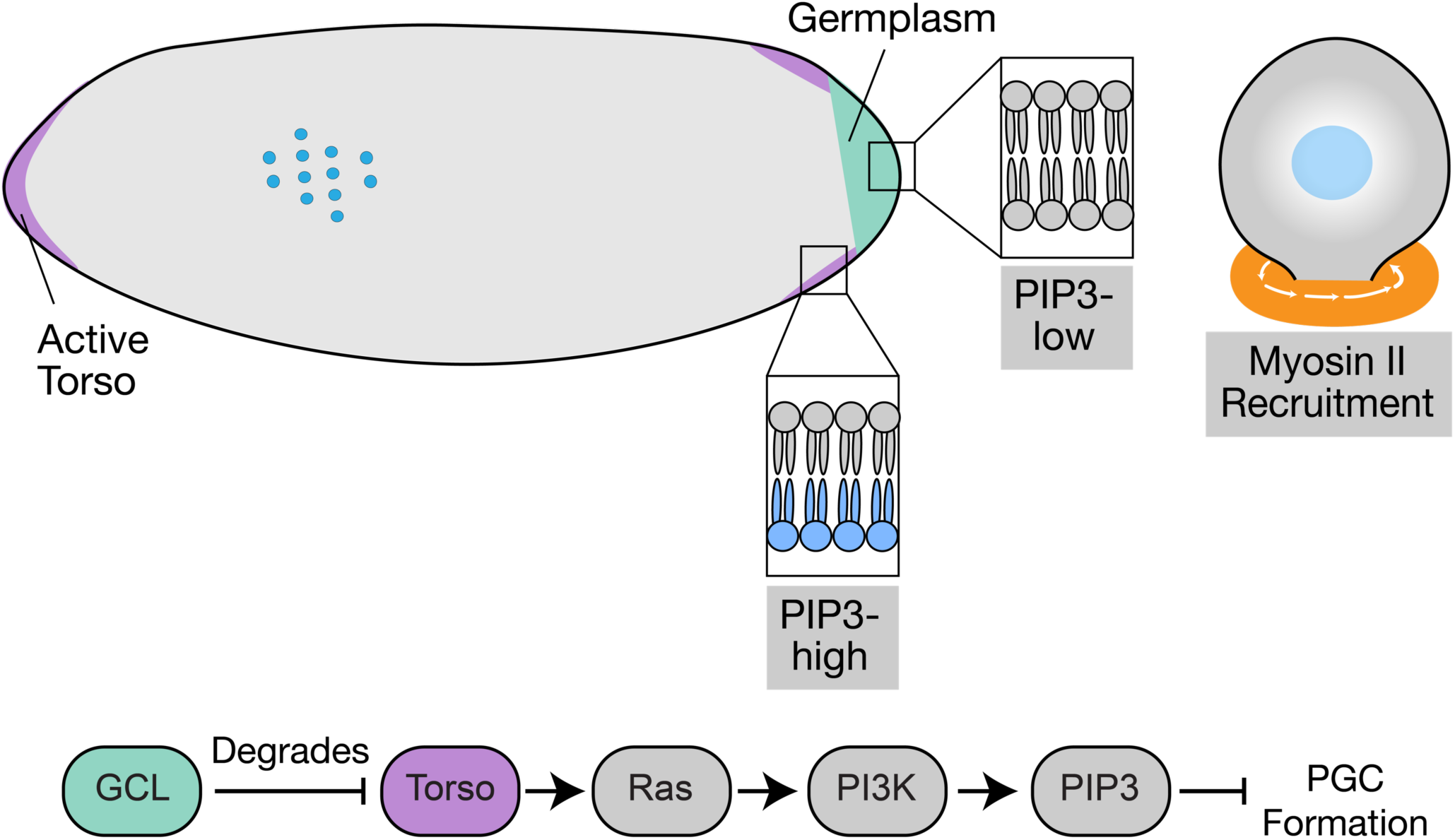
Graphical Abstract. - Membrane domains with high Torso activity (purple) in the early embryo have a higher PIP3 content.
- At the posterior pole, GCL-containing germplasm (green) degrades Torso, resulting in a PIP3-low membrane.
- Clearing PIP3 enables Myosin II pole bud constriction required for PGC formation.

## Introduction

Most embryos develop from a large egg that, upon fertilization, is divided into cells. In most insects, cellularization is delayed, allowing the embryo to initiate embryogenesis in a syncytial phase, with nuclei dividing within a shared cytoplasm. This process has been studied most extensively in *Drosophila melanogaster* where, after fertilization, nuclei undergo rapid synchronous divisions. After nine divisions, the nuclei migrate to the embryo’s cortex, where membrane remodeling leads to the formation of pseudocleavage furrows that bulge from the cortex (Foe and Alberts 1983). Only at the posterior pole of the embryo do these buds form cells, while the remainder of the embryo continues syncytial divisions. This difference arises from the posterior enrichment of a specialized cytoplasmic compartment, the germplasm, which the mother deposits into the embryo during oogenesis. The maternally contributed germplasm contains all the factors necessary to form and specify the embryo’s first cells, pole cells or primordial germ cells (PGCs), which will develop into the germline in the resulting adult (Illmensee and Mahowald 1974). Zygotic transcription is not essential for the initial formation of PGCs, making PGC formation an attractive model for studying early cell formation in other organisms that also depend on maternally provided factors (Cinalli and Lehmann 2013). In contrast, somatic cells form after the 13th nuclear division in a specialized membrane-deposition process that requires new zygotic transcription by the embryo.

PGC formation occurs through two orthogonal constrictions: a “traditional” spindle-dependent anaphase furrow constriction that divides one pole bud into two, and a spindle-independent “bud furrow” constriction that pinches off the dividing pole buds from the rest of the embryo cytoplasm (Cinalli and Lehmann 2013). The constriction of these two bud furrows must happen within a brief window during nuclear cycle 10 for PGCs to fully detach from the embryo, thereby protecting the integrity of the PGC-specifying germplasm (Lerit and Gavis 2011). How these two constrictions are regulated and coordinated remains unclear. After both furrows constrict and PGCs form, germplasm and nuclei are entirely segregated from the rest of the soma, allowing PGC specification to proceed. Once encased in a membrane, PGCs divide independently of the syncytial nuclear divisions. They undergo up to two divisions until the end of nuclear cycle 14, when morphogenetic gastrulation movements initiate their migratory journey through the embryo toward the somatic gonad (Chen et al. 2025)

One component of germplasm, Germ Cell-Less (GCL), is critical for the formation of PGCs (Jongens et al. 1994). Without maternally deposited *gcl* RNA and its localized translation at the posterior pole, *gcl-/-* embryos (referring to the maternal genotype) fail to undergo bud furrow constriction by the end of nuclear cycle 10, although anaphase furrow constrictions continue. As a result, pole buds fail to cellularize, and because germplasm is not properly sequestered and protected in PGCs, it is degraded. A minority of *gcl-/-* embryos (less than 20%) will form PGCs, though with greatly reduced numbers. GCL encodes a ubiquitin ligase adaptor, and the receptor tyrosine kinase (RTK) Torso was identified as its specific substrate for degradation (Pae et al. 2017). Torso’s RTK activity is restricted to the anterior and posterior regions of the developing oocyte due to the spatially regulated processing of its ligand, Trunk (Sprenger and Nüsslein-Volhard 1992; Stevens et al. 1990). Through GCL-mediated degradation of Torso, RTK activity is absent from the pole buds at the posterior pole. Indeed, Torso’s depletion at the posterior pole is necessary for PGCs to form, as the *gcl-/-* PGC defect can be fully rescued by preventing Torso’s RTK activity (Pae et al. 2017). Therefore, GCL shields posterior pole buds from RTK interference as they cellularize, allowing a parallel pathway, regulated by Rho1, anillin, and other components of the contractile ring, to direct pole bud cellularization (Field et al. 2005; Padash Barmchi et al. 2005).

It remains unclear how Torso RTK antagonizes PGC formation. Torso’s signaling role in embryogenesis activates the Ras/Raf/MEK/ERK pathway and triggers the transcription of patterning genes, such as the *huckebein* and *tailless* transcription factors, which are necessary for developing the embryonic termini (Brönner and Jäckle 1991). However, several observations argue against a transcriptional role for Torso in PGC formation: First, the embryo has not fully activated its zygotic transcription and relies primarily on maternally deposited factors at the time of PGC formation. Second, if transcription needed to be suppressed to allow for PGC formation, one would expect that inhibiting RNA Polymerase II would rescue the *gcl-/-* phenotype, but it does not (Cinalli and Lehmann 2013). Finally, the *gcl-/-* mutant phenotype can be rescued by knocking down the ArfGEF Steppke (Lee et al. 2015), a regulator of the endocytosis pathway, suggesting that inappropriate activation of Torso affects cytoskeletal organization rather than transcription.

We investigated Torso signaling to identify downstream effectors that interfere with PGC formation. Our results demonstrate a role for Torso signaling at the onset of embryonic development, independent of its known, later function in the transcriptional activation of terminal patterning genes. We identify PI3K as a critical downstream component of Torso activation and reveal an unexpected role for this signaling pathway early in embryogenesis in defining the germline-soma boundary via lipid domains.

## Results

### Light-induced activation of Ras causes failure in PGC formation

Previous studies indicated that ectopic activation of the Torso RTK pathway represses the cellular processes leading to PGC formation (Pae et al. 2017). As the canonical Torso RTK pathway recruits adaptor molecules to the membrane that activate Ras (Li 2005), we aimed to determine whether Ras activation is sufficient to interfere with PGC formation. Therefore, we employed an optogenetic approach, Optosos, which allows for precise spatiotemporal activation of Ras independent of RTK activity (Johnson et al. 2017). This is achieved by blue light-induced translocation of Sos, a Ras guanine nucleotide exchange factor (GEF), to the plasma membrane (OptoSos) (Toettcher et al. 2013; Johnson et al. 2017). This precise spatiotemporal control of Ras activation has two major advantages: (1) we could activate Ras specifically at the posterior pole while allowing the rest of the embryo to develop normally; and (2) we could induce ectopic signaling at a developmental stage before the Torso receptor activates embryonic somatic patterning (Casanova and Struhl 1989). Thus, instead of Ras activation at the blastoderm stage when PGCs had already formed (Johnson et al. 2017), we exposed the posterior pole of embryos to blue light at the syncytial stage prior to PGC formation (starting at 40 mins after egg deposition) (Fig 1A). To visualize the developmental process during time-lapse imaging, we also expressed the nuclear marker His2AV:RFP. Live imaging showed that pole buds still formed and protruded from the embryo cortex, but PGCs largely failed to cellularize, while control embryos that lacked the OptoSos blue light sensor formed PGCs normally (Fig 1B).

**Figure 1:**
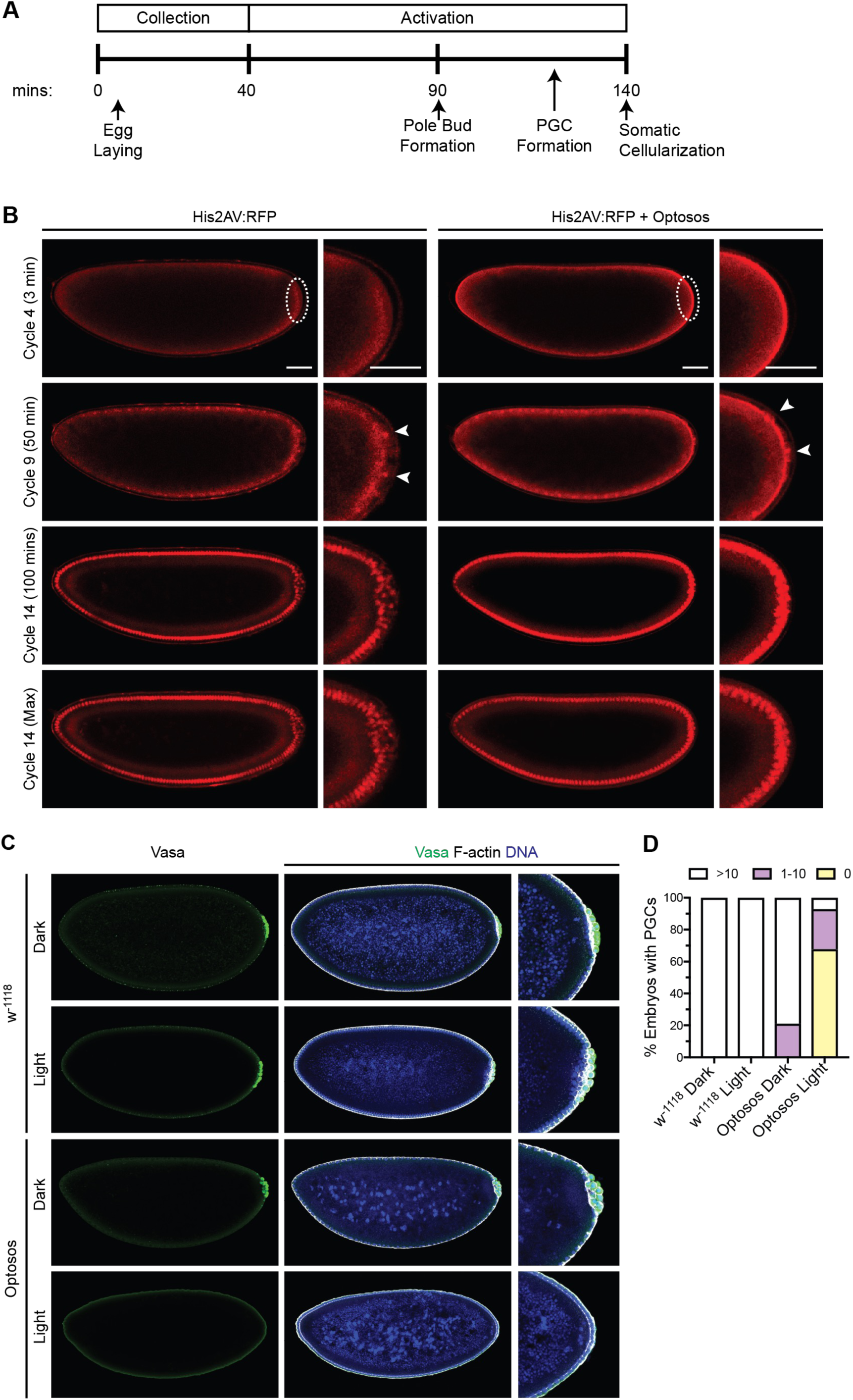
The RasGEF activity of Sos is sufficient to inhibit PGC formation, but not pole bud formation. (A) Timeline for optogenetic Ras activation. (B) Embryos expressing His2AV:RFP alone or His2AV:RFP and the OptoSos construct together were imaged with blue light activation in a region of interest (outlined with a dotted white oval). Arrowheads indicate the pole bud nuclei that protrude against the posterior membrane. In embryos that express His2AV:RFP alone, a group of rounded nuclei was observed adjacent to the layer of elongated somatic nuclei. This group of rounded nuclei was not seen in embryos that express both His2AV:RFP and the OptoSos construct. First three rows depict single z-plane images, while bottom row depicts a maximum intensity projection. Scale bar = 50μm. (C) Wild-type (w^-1118^) or OptoSos embryos were either kept in dark (Dark) or activated with blue light at the posterior pole (Light). After the 100-minute-long cycle of activation, embryos were fixed and stained with an antibody against Vasa to assess PGC formation. Percentage of embryos with the indicated number of PGCs was scored. (D) Representative embryo for each indicated condition. Fixed embryos were immunostained with anti-Vasa (Green). F-actin (gray) outlines the membrane. DNA (blue).

To quantitatively assess these live observations, embryos were immunostained with an antibody against Vasa to count PGCs at nuclear cycle 14, when PGC divisions had ceased (Fig 1C). Ninety percent of embryos expressing OptoSos that were activated with blue light during the syncytial stage showed a reduction or complete loss of PGCs compared to control embryos that were kept in the dark (20% showed a reduced number of PGCs) or w^-1118^ embryos that received either dark or light treatment (no reduction in PGCs) (Fig 1D). These data demonstrate that early activation of Ras is sufficient to prevent pole buds from completing cellularizing, even in the presence of GCL.

### Torso activation interferes with PGC formation independently of the Raf-MEK-ERK pathway

Since the optogenetic induction of Ras was sufficient to block PGC formation, we next examined which effectors downstream of Ras were necessary for this process. Since genetic disruption of Torso RTK restored PGC formation in *gcl-/-* embryos (Pae et al. 2017), we conducted RNAi against downstream components of the Torso pathway in *gcl-/-* mutant embryos and assessed whether PGC formation was similarly restored (Fig. 2A-C). As expected, based on genetic and optogenetic experiments, RNAi knockdown of *torso*, *shc*, *sos*, and *ras* restored PGC formation in *gcl-/-* embryos (Fig 2B). However, knockdown of *csw* (the fly homolog of SHP2), *raf*, *ksr*, *dsor1* (the fly homolog of MEK), and *rolled* (the fly homolog of MAPK) did not restore PGC formation (Pae et al. 2017). To verify that RNAi effectively disrupted Torso signaling under these conditions, we stained for dpERK, a marker for MAPK phosphorylation, the final kinase step in canonical RTK signaling (Fig. 2C). All RNAi conditions except for mCherry RNAi displayed a loss of dpERK staining at the anterior and posterior poles. These results demonstrate that the activation of the canonical RTK pathway downstream of Ras, including the Raf/MEK/MAPK signaling cascade, which plays a critical role in the transcriptional activation of somatic target genes, is neither necessary nor sufficient to interfere with PGC formation. We conclude that another, previously unknown effector of Torso RTK is involved in PGC formation.

**Figure 2:**
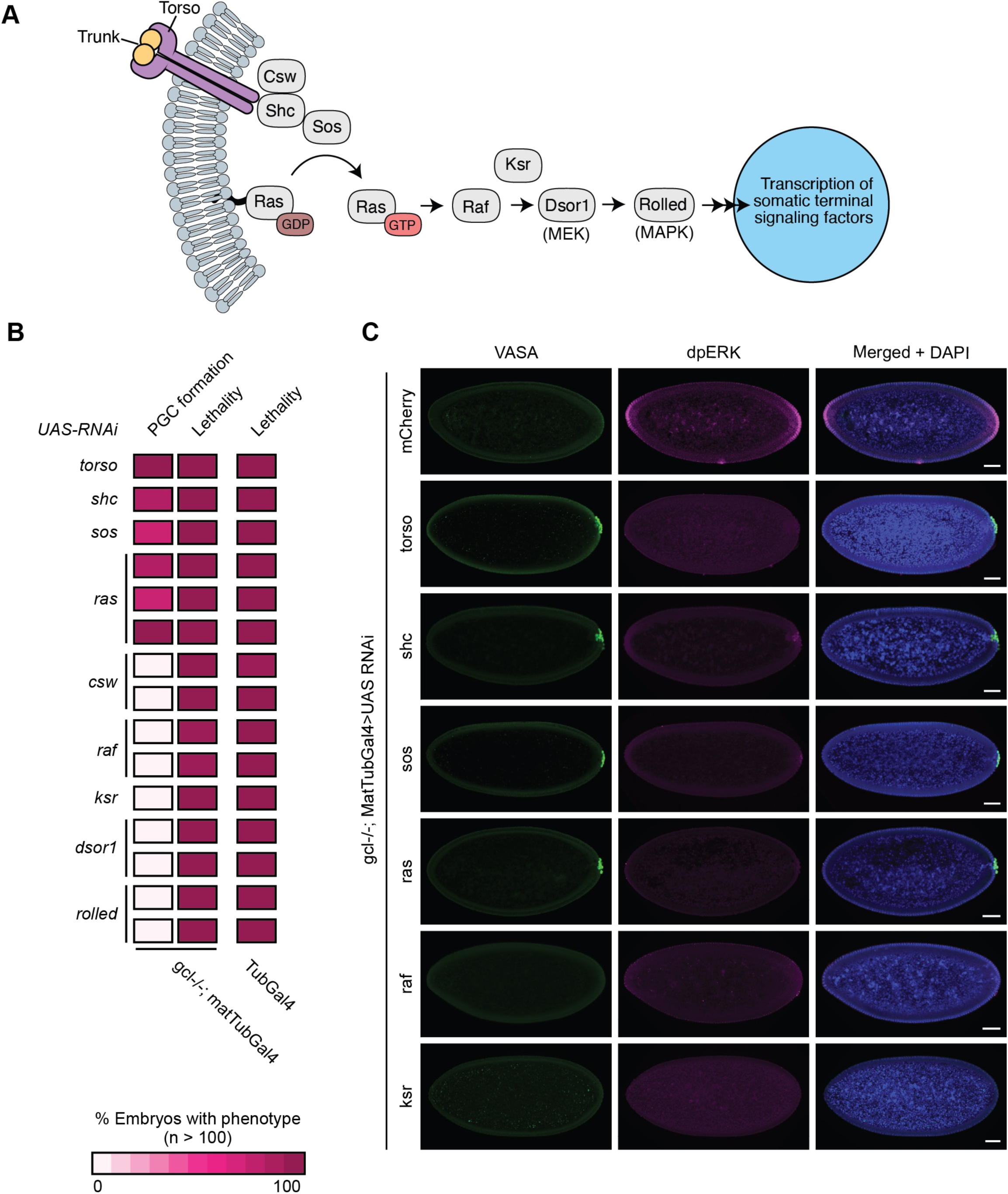
Ras is the most downstream component in the canonical Torso signaling pathway that is involved in antagonizing PGC formation. (A) Diagram of the canonical Torso signaling pathway which is required to upregulate transcription of somatic patterning genes in the embryo (B) The canonical Torso signaling pathway components were knocked down in *gcl-/-* embryos using RNAi driven with the germline-specific driver matTubGal4. Percentage of embryos with Vasa-positive PGCs was calculated and plotted as a score for PGC formation. Lethality in these embryos was also scored. The ubiquitous driver TubGAL4 was used to assess knockdown efficiency (n>100). (C) Representative embryo of indicated genotypes. Fixed embryos were immunostained with anti-Vasa (Green) and anti-dpERK (magenta). DNA (blue). Scale bar = 50μm.

### Ras separation-of-function mutants suggest that PI3K or RalA activation interfere with PGC pathway activation

To identify the signaling pathway downstream of Ras that may affect PGC formation, we focused on two additional, known downstream effectors of Ras: RalA and PI3K. The functions of these signaling pathways can be analyzed independently of the canonical MAP kinase pathway using a separation-of-function strategy. Point mutations in the effector loop domain of Ras act as molecular switches that, in addition to the Ras-V12 mutation, allow for the constitutive activation of a specific downstream pathway, while blocking Ras’s ability to activate other downstream pathways (Rodriguez-Viciana et al. 1997, Prober and Edgar 2002). For example, the Ras-S35 mutation enables the constitutive activation of the Raf pathway while inhibiting Ras’s ability to activate RalA and PI3K (Fig. 3C). To specifically separate Ras’s ability to activate its effectors, we knocked down endogenous *ras* expression with RNAi and expressed the effector loop mutants from RNAi-insensitive constructs (Fig 3A). Interestingly, knockdown of *ras* alone increased the number of PGCs by 30% compared to control embryos, demonstrating that Ras signaling counteracts PGC formation even in wild-type embryos, when GCL is present (Fig. 3D).

**Figure 3:**
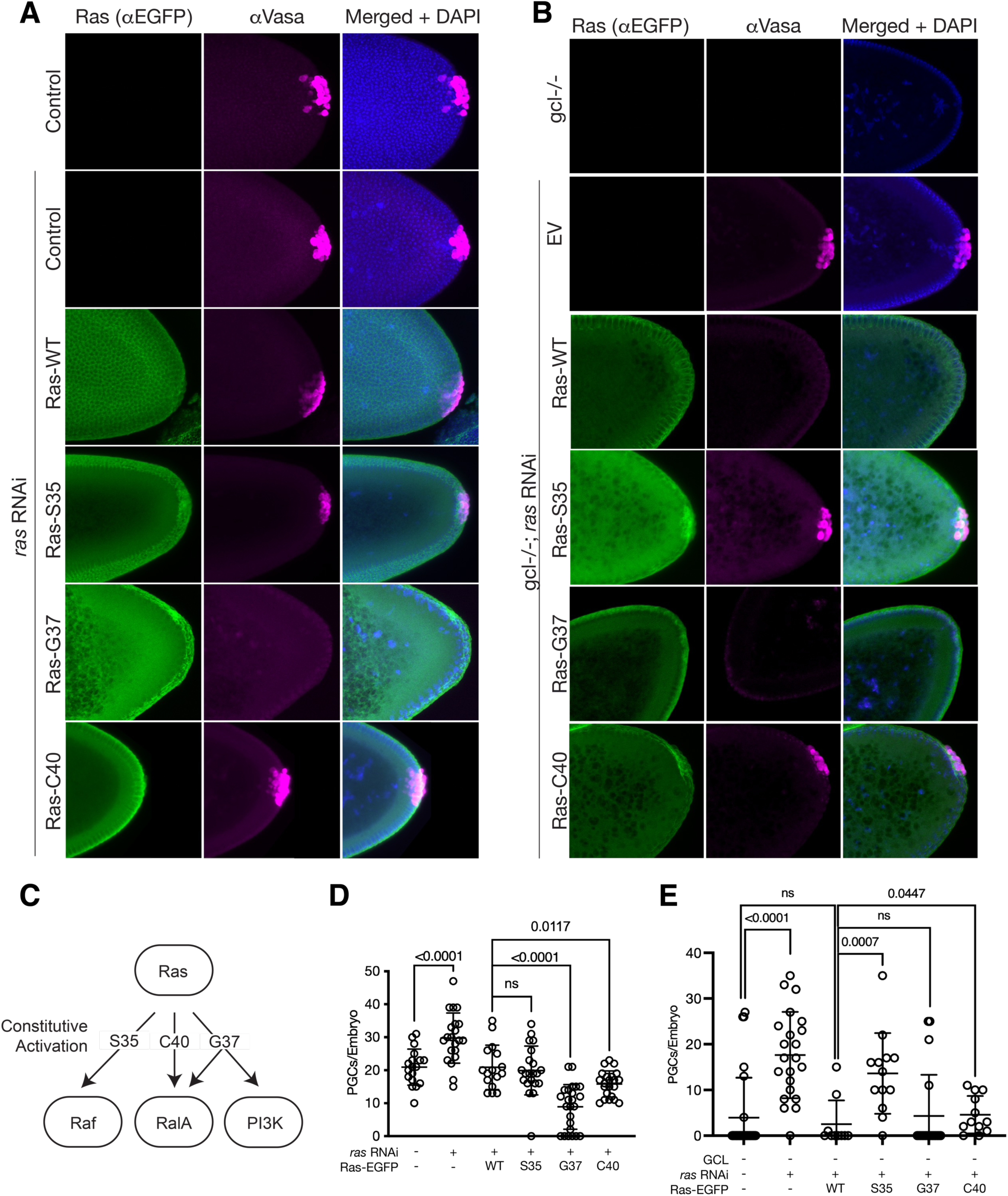
Expression of Ras effector loop mutants that activate PI3K decreases PGC formation, while Raf activating mutants do not affect PGC formation. (A) Embryos expressing Ras RNAi and RNAi-insensitive EGFP-tagged Ras, Ras-S35, Ras-G37, and Ras-C40 were immunostained with anti-EGFP and anti-VASA to confirm expression of the construct and stain PGCs for counting. Expression of the UASp constructs was driven with two copies of maternal-tubulin-GAL4-VP16 for optimal expression during late oogenesis. Images depict maximum intensity projections spanning area of PGC formation. (B) *gcl-/-* embryos expressing Ras RNAi and RNAi-insensitive EGFP-tagged Ras, Ras-S35, Ras-G37, and Ras-C40 were immunostained with anti-EGFP and anti-VASA to confirm expression of the construct and stain PGCs for counting. Expression of the UASp constructs was driven with maternal-tubulin-GAL4-VP16 for optimal expression during late oogenesis. Images depict maximum intensity projections spanning the area of PGC formation (C) Model illustrating downstream pathways that are constitutively activated by Ras effector loop mutations. (D) PGCs in nuclear cycle 13-14 embryos from mothers of the indicated genotype were counted and plotted from (A). The maternal genotype of control embryos is w1118. Bars represent the mean ± standard deviation. (n > 15, Mann-Whitney test) (E) PGCs in nuclear cycle 13-14 embryos from mothers of the indicated genotype were counted and plotted from (B). The maternal genotype of control embryos is w1118. Bars represent the mean ± standard deviation. (n > 10, Mann-Whitney test)

Overexpression of wild-type Ras restored the number of PGCs to control levels. Embryos overexpressing Ras-S35 did not alter the number of PGCs formed, while embryos expressing Ras-G37 and Ras-C40 showed a reduction in PGCs, with a 57.5% reduction in Ras-G37 embryos (Fig. 3D). Since Ras-G37 is known to activate both the RalA and the PI3K pathways, and Ras-C40 activates the PI3K pathway, our results suggest that these alternative pathways downstream of Ras interfere with PGC formation (Pacold et al. 2000; Kinashi et al. 2000).

Since Ras effector mutants are primarily thought to activate only one branch of the signaling pathway while inhibiting the other pathways, we used the separation-of-mutants to test which Ras effectors are targeted by GCL function. We expressed the Ras effector loop mutants in *gcl-/-* embryos alongside *ras* RNAi and measured PGC formation (Fig 3B). The expression of Ras-S35 rescued PGC formation in *gcl-/-* embryos, further demonstrating that the Ras-Raf pathway does not cause the PGC defect. The Ras-S35 mutant can no longer activate other downstream pathways of Ras, such as RalA and PI3K, thus blocking Ras’ ability to antagonize PGC formation. Overexpression of Ras-G37 could not rescue PGC formation in *gcl-/-* embryos, while overexpression of Ras-C40 was able to partially rescue PGC formation, though not to the same degree as Ras-S35 (Fig. 3E). These results support the hypothesis that the activation of RalA and PI3K pathways antagonizes PGC formation and that this activation contributes to the PGC formation defects in *gcl-/-* embryos.

### PI3K activation is a major antagonist of PGC formation

Ras separation-of-function mutants identified two potential pathways downstream of Ras whose activation may affect PGC formation: RalA, a GTPase that recruits the exocyst complex and plays a role in membrane addition during somatic cellularization (Holly et al. 2015), and PI3K, a key regulator of metabolism and cellular morphogenesis which phosphorylates phosphatidylinositol 4,5-bisphosphate (PIP2) to generate phosphatidylinositol 3,4,5-trisphosphate (PIP3) (Stambolic et al. 1998; Shewan et al. 2011). To directly determine whether the activity of these effectors can interfere with PGC formation specifically, we expressed variants of RalA and PI3K subunits at the posterior pole using a germplasm-specific 3’UTR containing the translational control element of *nanos* and the 3’UTR of *pgc* (TCEp3) (Lin et al. 2020).

We generated constitutively active and dominant-negative mutations in RalA: RalA G20V and RalA-S25N, respectively. For human RalA, these mutations have been shown to modulate RalA activity by changing its binding affinity to GDP and GTP (Hinoi et al. 1996; Sawamoto et al. 1999). Germplasm-specific overexpression of RalA-G20V slightly decreased PGC formation by 21.7%, while germline overexpression of RalA-S25N, a dominant negative allele of RalA, had no effect (Fig. S1A-B). Similarly, germline overexpression of the RalGEF Rgl, which is expected to increase RalA activity, only slightly decreased PGC formation by 24.6% (Fig. S1C-D). In contrast, increasing PI3K activity had a robust antagonistic effect on PGC formation. PI3K activity relies on a catalytic subunit, p110 in mammals and dp110 in *Drosophila*, as well as a regulatory subunit, p85 in mammals and p60 in *Drosophila*. Germplasm overexpression of the wild-type catalytic domain dp110 significantly reduced PGC formation by 37.6% (Fig. 4A-B). Further increasing PI3K activity by overexpressing membrane-targeted dp110-CAAX had an even stronger effect, reducing PGC formation by 97.0% (Fig 4A-B). While our results suggest that activation of RalA and PI3K can counteract PGC formation, we observed more robust effects by manipulating the PI3K pathway. Indeed, while Ras-C40 and Ras-G37 have both been shown to activate PI3K signaling, it has previously been reported that expression of Ras-G37 but not Ras-C40 or Ras-S35 results in increased PIP3 levels in Drosophila wing tissues (Prober and Edgar 2002). We therefore focused our subsequent analysis on the mechanisms by which the PI3K pathway is regulated at the posterior pole and how this regulation affects PGC formation.

**Figure 4:**
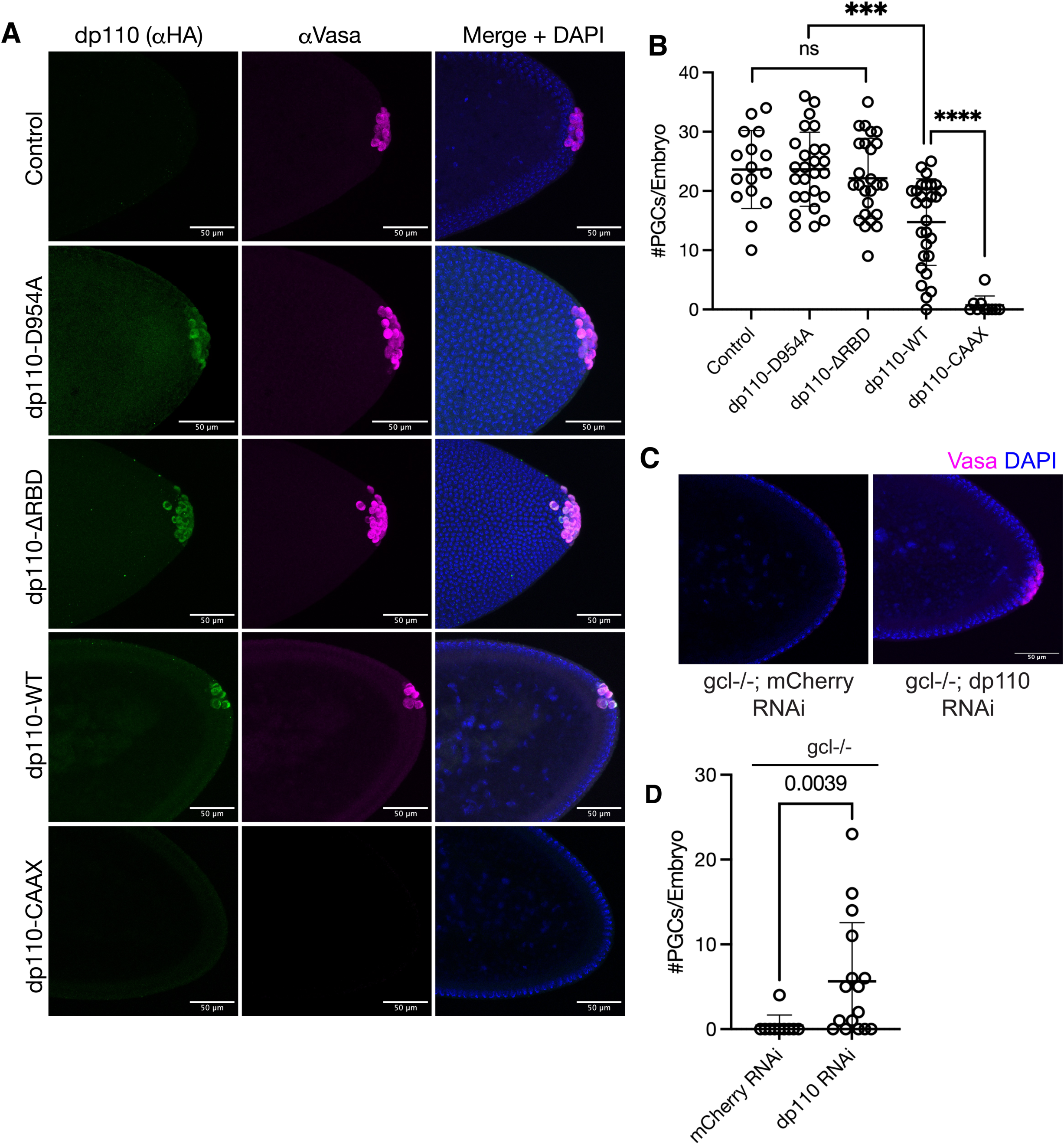
PI3K strongly antagonizes PGC formation through its Ras-binding and catalytic activity. (A) Embryos overexpressing germline-targeted, HA-tagged dp110, dp110-D954A (catalytically inactive), dp110-ΔRBD (Ras binding mutant), and dp110-CAAX (constitutively active) were immunostained with anti-HA and anti-VASA to confirm expression of the construct and stain PGCs for counting. Expression of the UASp constructs was driven with two copies of maternal-tubulin-GAL4-VP16 for optimal expression during late oogenesis. Images depict maximum intensity projections spanning area of PGC formation. Scale bar = 50 µm. (B) PGCs in nuclear cycle 13-14 embryos from mothers of indicated genotype were counted and plotted from (C). The maternal genotype of control embryos is maternal-tubulin-GAL4-VP16/+; maternal-tubulin-GAL4-VP16/+. Control, dp110-D954A, and dp110-ΔRBD embryos show no significant difference in PGC amounts. Bars represent the mean ± standard deviation. (n > 10, Mann-Whitney test, ***P < 0.001, ****P < 0.0001) (C) gcl-/- embryos expressing RNAi against *dp110* were immunostained with anti-Vasa to count PGCs. Images depict maximum intensity projections spanning area of PGC formation. Scale bar = 50 µm. (D) PGCs in nuclear cycle 13-14 embryos from mothers of indicated genotype were counted and plotted. RNAi against mCherry was used as a control. Bars represent the mean ± standard deviation. (n > 10, Mann-Whitney test)

To test whether dp110’s catalytic activity was required for its suppression of PGC formation, we overexpressed germ plasm-targeted dp110-D954A, which can no longer bind to ATP and has been shown to have a dominant negative phenotype when overexpressed (Leevers et al. 1996). These embryos formed the same amount of PGCs as wild-type control, suggesting that ATP-binding and catalytic activity is required for dp110 to suppress PGC formation (Fig 4A-B). To test whether the binding of the PI3K catalytic subunit to Ras mediated the effect on PGC formation, we mutated the Ras binding domain (RBD) of dp110. Four point mutations in the RBD have been shown to block dp110’s ability to bind to Ras and decrease the maximal signaling of PI3K, even though the catalytic domain remains intact (Orme et al. 2006). Consistent with a direct interaction between Ras and PI3K, dp110-ΔRBD overexpression did not affect PGC formation (Fig 4A-B).

Next, we inquired whether inappropriate PI3K activation could explain the defects observed in *gcl-/-* embryos. If so, we would expect that decreasing dp110 expression would restore PGC formation in *gcl*-/- embryos. Indeed, RNAi knockdown of the PI3K dp110 subunit in *gcl-/-* embryos partially rescued PGC formation, with 50% of embryos making five or more PGCs compared to 0% in control (Fig. 4C-D). These experiments identify PI3K as a direct mediator of Torso activation in antagonizing PGC formation.

### Suppression of PI3K activity results in ectopic cellularization only at the posterior pole

To determine how PI3K activation may interfere with PGC formation—a cellular event involving membrane budding and localized membrane constriction—we focused on the role of PI3K in converting PIP2 to PIP3. This conversion relies on the regulatory subunit of PI3K, p60, which functions as a switch that mediates the plasma membrane localization of dp110 upon Ras activation (Leevers et al. 1996). The pi3K21B gene encodes the p60 subunit in Drosophila, and the pi3k21b transcript is enriched in the germplasm of early embryos and in PGCs after their formation (Xi et al. 2014). Imaging of a sfGFP-tagged fosmid line revealed that p60 protein is present in high levels throughout the early embryo (S4A-B). P60 binds to dp110 and suppresses PI3K function in the absence of RTK activation (Leevers et al. 1996). Therefore, increasing p60 expression may further protect pole buds from PI3K activity. To test this idea, we overexpressed p60, which resulted in an increase in PGC formation compared to the control, demonstrating the ability of p60 in suppressing PI3K activity during PGC formation (Fig. 5A-B). These constructs do not contain the germline-targeting hybrid 3’UTR and express p60 uniformly across the embryo (Tamada et al. 2021). Despite this, there was no evidence of ectopic cellularization of the soma during nuclear cycle 14 when p60 was overexpressed, indicating that this effect is specific to the posterior pole where germplasm is enriched. The SH2 domains of PI3K’s regulatory subunit suppress PI3K activity, and removal of the SH2 domains has been shown to increase PI3K activity (Klippel et al. 1994). When we overexpressed p60-ΔSH2, we observed a decrease in PGC formation similar to membrane-targeted dp110, supporting the conclusion that p60 can act as a modulator of PI3K signaling that is particularly critical for PGC formation (Fig. 5A-B).

**Figure 5:**
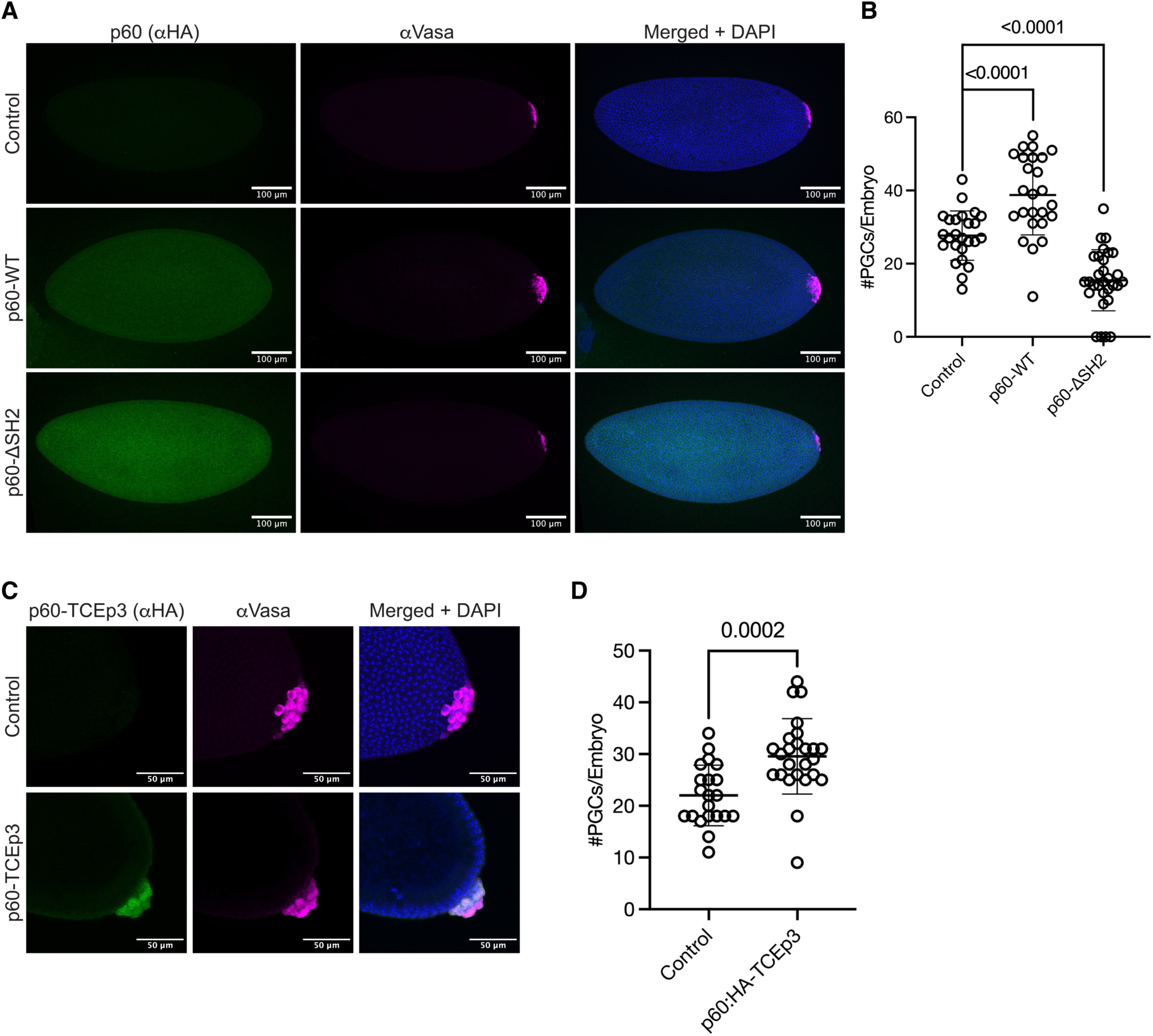
Decreasing PI3K activity rescues the *gcl* mutant phenotype and increases pole cell formation. (A) Embryos overexpressing HA-tagged p60 and p60-ΔSH2 were immunostained with anti-HA and anti-Vasa to confirm expression and count PGCs. Images depict maximum intensity projections spanning area of PGC formation. Scale bar = 100 µm. (B) PGCs in nuclear cycle 13-14 embryos from mothers of indicated genotype were counted and plotted. The maternal genotype of control embryos is maternal-tubulin-GAL4-VP16/+; maternal-tubulin-GAL4-VP16/+. Bars represent the mean ± standard deviation. (n > 20, Mann-Whitney test) (C) Embryos overexpressing germplasm-targeted HA-tagged p60 were immunostained with anti-HA and anti-Vasa to confirm expression and count PGCs. Images depict maximum intensity projections spanning area of PGC formation. Scale bar = 50 µm. (D) PGCs in nuclear cycle 13-14 embryos from mothers of indicated genotype were counted and plotted. The maternal genotype of control embryos is maternal-tubulin-GAL4-VP16/+; maternal-tubulin-GAL4-VP16/+. Bars represent the mean ± standard deviation. (n > 20, Mann-Whitney test).

Germplasm-targeted overexpression of wild-type p60 corroborated this result by increasing PGC formation compared to the control. In addition, PGCs formed in these embryos exhibited a hyper-clustered phenotype, where they protruded from the embryo’s somatic blastoderm and tightly packed on top of each other (Fig. 5C-D). This contrasts with control embryos, where PGCs form a less compact monolayer that wraps around the posterior pole. Our results clearly demonstrate a connection between PI3K activity and PGC formation: low levels of PI3K activity support PGC formation, whereas high levels antagonize it.

### Spatially restricted Torso activity patterns PIP3 in the embryo

PI3K catalyzes the production of PIP3 by phosphorylating PIP2. To understand how this production is impacted by GCL-mediated degradation of Torso at the posterior pole, we analyzed the distribution of PIP3 in wild-type and *gcl-/-* embryos using the biosensor GFP:Grp1[PH] (Prober and Edgar 2002). In early wild-type embryos, before nuclei have reached the embryo cortex, PIP3 was enriched at the membrane in areas of high Torso activity: the anterior pole and the regions adjacent to the posterior pole, but was excluded from the posterior pole where germplasm is localized (Fig. 6A). In *gcl-/-* embryos, PIP3 was no longer excluded from the posterior pole. Thus, by promoting degradation of the Torso RTK, GCL prevents PIP3 generation specifically at the posterior pole where it is translated.

**Figure 6:**
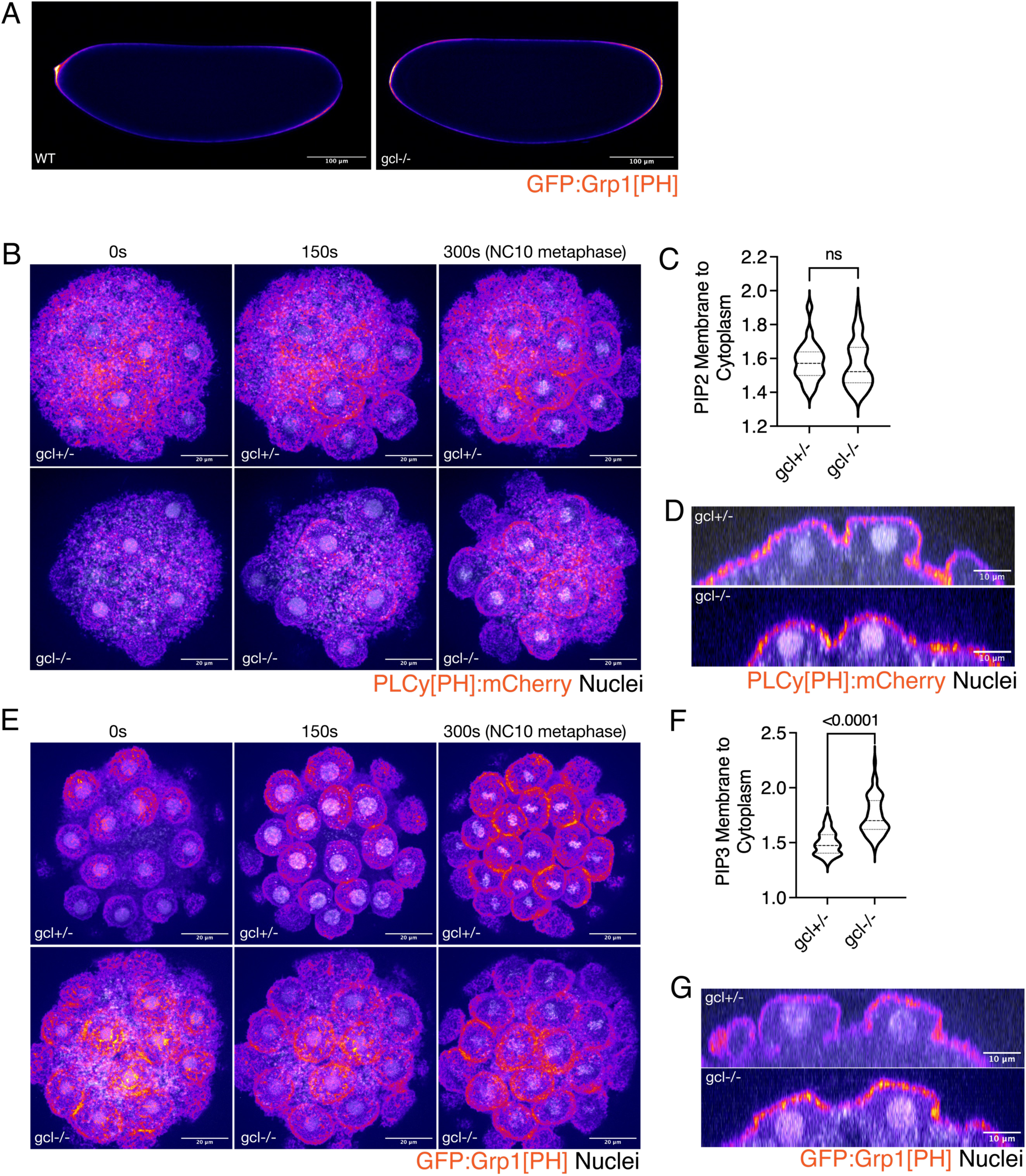
PIP3 acts downstream of Torso, and is inappropriately enriched in the posterior pole membrane when GCL is not present. (A) Wild-type and *gcl-/-* embryos expressing the PIP3 biosensor GFP:Grp1[PH] (fire) were live imaged on their side to visualize PIP3 spatial distribution. Embryos are less than an hour old, since nuclei have not yet migrated to cortex. Scale bar = 100 µm. Embryos are oriented with their anterior poles to the left and their posterior poles to the right. Images show single plane roughly in the middle of the embryo. Scale bar = 100 µm. (B) *gcl+/-* and *gcl-/-* nuclear cycle 10 embryos expressing the PIP2 biosensor PLCγ[PH]:mCherry (fire) and nuclear marker His2AV:GFP (gray) were posteriorly mounted and live imaged. Images are maximum intensity projections of a 20 µm section. The 300s time point represents nuclear cycle 10 metaphase. Scale bar = 20 µm. (C) Membrane and cytoplasm were segmented from posterior-mounted embryo movies 150s before nuclear cycle 10 metaphase, and the membrane-to-cytoplasm ratio of PIP2 fluorescence was plotted. (n = 6 embryos for gcl+/-, n = 8 embryos for gcl-/-, Mann-Whitney test) (D) Orthogonal view of *gcl +/-* and *gcl-/-* embryos expressing the PIP2 biosensor PLCγ[PH]:mCherry (fire) and nuclear marker His2AV:GFP (gray) 150s before nuclear cycle 10 metaphase. Scale bar = 10 µm. (E) *gcl+/-* and *gcl-/-* nuclear cycle 10 embryos expressing the PIP3 biosensor GFP:Grp1[PH] (fire) and nuclear marker His2AV:RFP (gray) were posteriorly mounted and live imaged. Images are maximum intensity projections of a 20 µm section. The 300s time point represents nuclear cycle 10 metaphase. Scale bar = 20 µm. (F) Membrane and cytoplasm were segmented from posterior-mounted embryo movies 150s before nuclear cycle 10 metaphase, and the membrane-to-cytoplasm ratio of PIP3 fluorescence was plotted. (n = 6 embryos for gc*l+/-,* n = 5 embryos for *gcl-/-,* Mann-Whitney test) (G) Orthogonal view of *gcl +/-* and *gcl-/-* embryos expressing the PIP3 biosensor GFP:Grp1[PH] (fire) and nuclear marker His2AV:RFP (gray) 150s before nuclear cycle 10 metaphase. Scale bar = 10 µm

PIP2 has recently been shown to be enriched at the posterior pole in the cellularizing pole buds (Kilwein et al. 2025). We used the PIP2 biosensor PLCy[PH]:mCherry and the PIP3 biosensor GFP:Grp1[PH] to simultaneously observe the distribution of these phosphoinositide species in developing embryos. While PIP3 is restricted to membrane regions with high Torso activity (Fig. 6A), PIP2 exhibits a more dynamic localization, enriching at either the anterior or posterior pole depending on the movement of the entire membrane in the cell cycle during syncytial divisions (Fig. S1A, Movie 1). Examining the dynamic localization of these phosphoinositides in pole buds by live imaging, we observed that PIP3 is depleted from the base of pole buds, while PIP2 is enriched at the base and throughout the bud membrane (Fig S2B-C). To measure if there was a change in PIP2 and PIP3 levels in the membrane of *gcl-/-* pole buds, we live-imaged either PLCy[PH]:mCherry or GFP:Grp1[PH] in posteriorly mounted embryos along with the nuclear marker His2AV:GFP or His2AV:RFP. This enabled precise embryo staging and pole bud segmentation, thereby capturing membrane dynamics during nuclear cycle 10 immediately preceding PGC formation. The membrane enrichment of PIP3, as measured by the membrane-to-cytoplasm ratio of PIP3 fluorescence intensity, is higher in *gcl-/-* nuclear cycle 10 posterior pole buds compared to *gcl+/-* pole buds, while there is no difference in PIP2 fluorescence intensity (Fig. 6B-G). This suggests that the previously reported posterior enrichment of PIP2 is independent of GCL’s function (Kilwein et al. 2025). This also aligns with previous observations that the PI3K pathway does not significantly affect PIP2 levels (Jackson et al. 1992).

To test whether loss of Torso signaling abolishes the spatial patterning of PIP3, we imaged GFP:Grp1[PH] in loss-of-function *torso^HH/WK^*embryos (Fig. 7A-B). While we no longer observed a wild-type pattern of PIP3, the biosensor appeared to enrich to the anterior and posterior pole in early embryos. We tested whether Torso-like, which can have roles independent of Torso RTK signaling, may contribute to the terminally enriched PIP3 patterning (Taylor et al. 2019). We imaged GFP:Grp1[PH] in embryos from *tsl^3/4^*mothers and observed that the terminal enrichment of PIP3 persisted as in *torso* mutant embryos (Fig. 7A). We hypothesize that in the absence of PIP3 production, the strongly expressed biosensor may bind promiscuously to other phosphoinositides, since the PH domain of Grp1 has been shown to have some nonspecific binding (Li et al. 2020). To more directly determine the sensitivity of PIP3 levels to Torso activity at the time of PGC formation, we compared PIP3 membrane enrichment in embryos with various levels of posterior pole Torso activity. As expected for GCL-dependent degradation of Torso, the membrane-to-cytoplasm ratio of GFP:Grp1[PH] was indistinguishable between wild-type and *torso^HH/WK^* pole buds at 150 seconds before nuclear cycle 10 (Fig. 7B-C). Lower levels of PIP3 were observed at the onset of metaphase at the 300s time point in *torso^HH/WK^*pole buds, indicating that wild-type posterior pole buds have latent levels of Torso and PI3K signaling that escape GCL-mediated degradation (Fig. 7D). PIP3 levels were significantly elevated in *gcl*-/- pole buds compared to wild-type. Interestingly, we also observed higher PIP3 levels in *gcl*+/- pole buds, demonstrating the sensitivity of PIP3 to Torso activity (Fig. 7C). This also suggests there is a threshold level of PIP3 that can be tolerated for PGC formation, as *gcl+/-* embryos produce approximately the same number of PGCs as wild-type.

**Figure 7:**
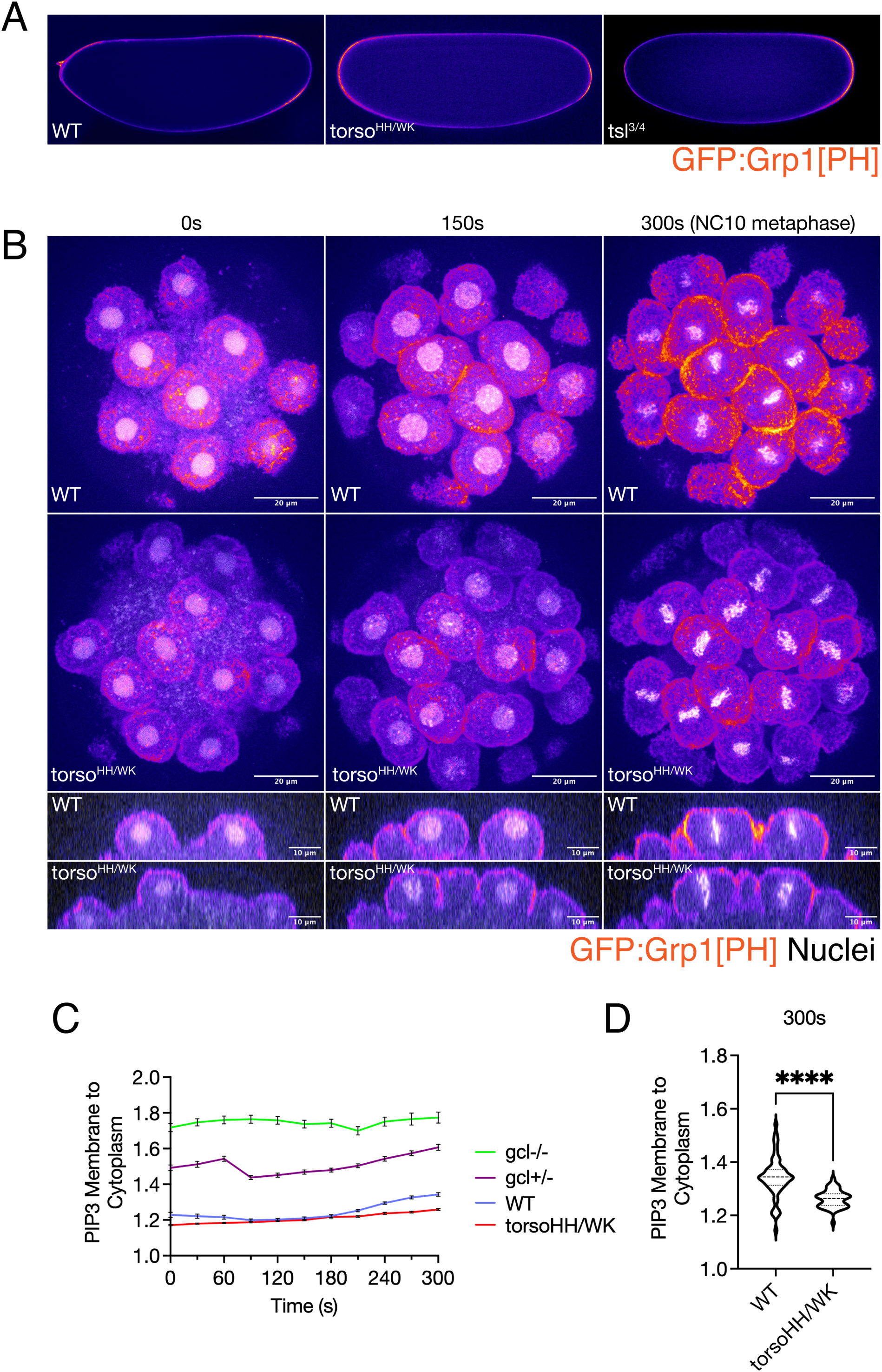
PIP3 levels in the embryo are dependent on the dosage of Torso activity. (A) Wild-type, *torso^HH/WK^*, and *tsl^3/4^* embryos expressing the PIP3 biosensor GFP:Grp1[PH] (fire) were live imaged on their side to visualize PIP3 spatial distribution. Embryos are less than an hour old, since nuclei have not yet migrated to cortex. Embryos are oriented with their anterior poles to the left and their posterior poles to the right. Images show single plane roughly in the middle of the embryo. (B) Wild-type and torso HH/WK nuclear cycle 10 embryos expressing the PIP3 biosensor GFP:Grp1[PH] (fire) and nuclear marker His2AV:RFP (gray) were posteriorly mounted and live imaged. Images are maximum intensity projections of a 20 µm section. The 300s time point represents nuclear cycle 10 metaphase. Scale bar = 20 µm. Lower panels show orthogonal views. Scale bar = 10 µm. (C) Timelapse of PIP3 membrane to cytoplasm in wild-type, *gcl+/-*, *gcl-/-,* and *torso^HH/WK^* embryos. Bars represent the mean ± SEM. (D) The membrane to cytoplasm ratio of PIP3 fluorescence at nuclear cycle 10 metaphase (300s timepoint) was plotted for wild-type and torso HH/WK embryos. (n = 7 embryos for wild-type, n = 7 embryos for torso HH/WK, Mann-Whitney test)

Together, these results suggest that the spatially restricted activation of Torso at the egg termini patterns PI3K activity and PIP3 enrichment in the early embryo. GCL-dependent degradation of Torso at the posterior pole causes asymmetrical membrane remodeling at the site of PGC formation.

Pole buds in *gcl-/-* embryos show defective myosin assembly.

Cellular morphogenesis events such as cytokinesis and cell migration depend on precise spatiotemporal regulation of PIP2 and PIP3 to effectively balance protrusive and contractile forces (Shewan et al. 2011). During *Drosophila* somatic cellularization, excessive PIP3 results in failed cellularization due to myosin disassembly, while excessive PIP2 causes premature cellularization due to increased contractility (Reversi et al. 2014). The architecture of the round, protrusive pole bud relies significantly on the counterforces between the cortical stiffness provided by the F-actin-rich cap and the contractility generated by the actomyosin-rich base (Fernandez-Gonzalez and Harris 2023). Elevated PIP3 levels often indicate rapid actin polymerization, particularly at the leading edges of migrating cells (Devreotes and Janetopoulos 2003). To investigate whether PIP3 levels correlated with F-actin polymerization in the caps of pole buds during PGC formation, we live-imaged the actin-binding domain of moesin fused to GFP (GFP:moe[ABD]). GFP:moe[ABD] serves as a reliable marker of F-actin that does not interfere with endogenous actin dynamics (Spracklen et al. 2014). Our findings revealed that in *gcl-/-* embryo nuclear cycle 10 pole buds, F-actin appeared disorganized and roughened, but the fluorescence intensity did not increase when compared to *gcl+/-* pole buds, suggesting that actin polymerization did not increase (Fig. 8A-C). The roughened appearance of the F-actin caps could be a result of excessive membrane ruffling, which is induced by growth factor stimulation of PI3K and PIP3 (Rodriguez-Viciana et al. 1997; Deming et al. 2008; Yan et al. 2024).

**Figure 8:**
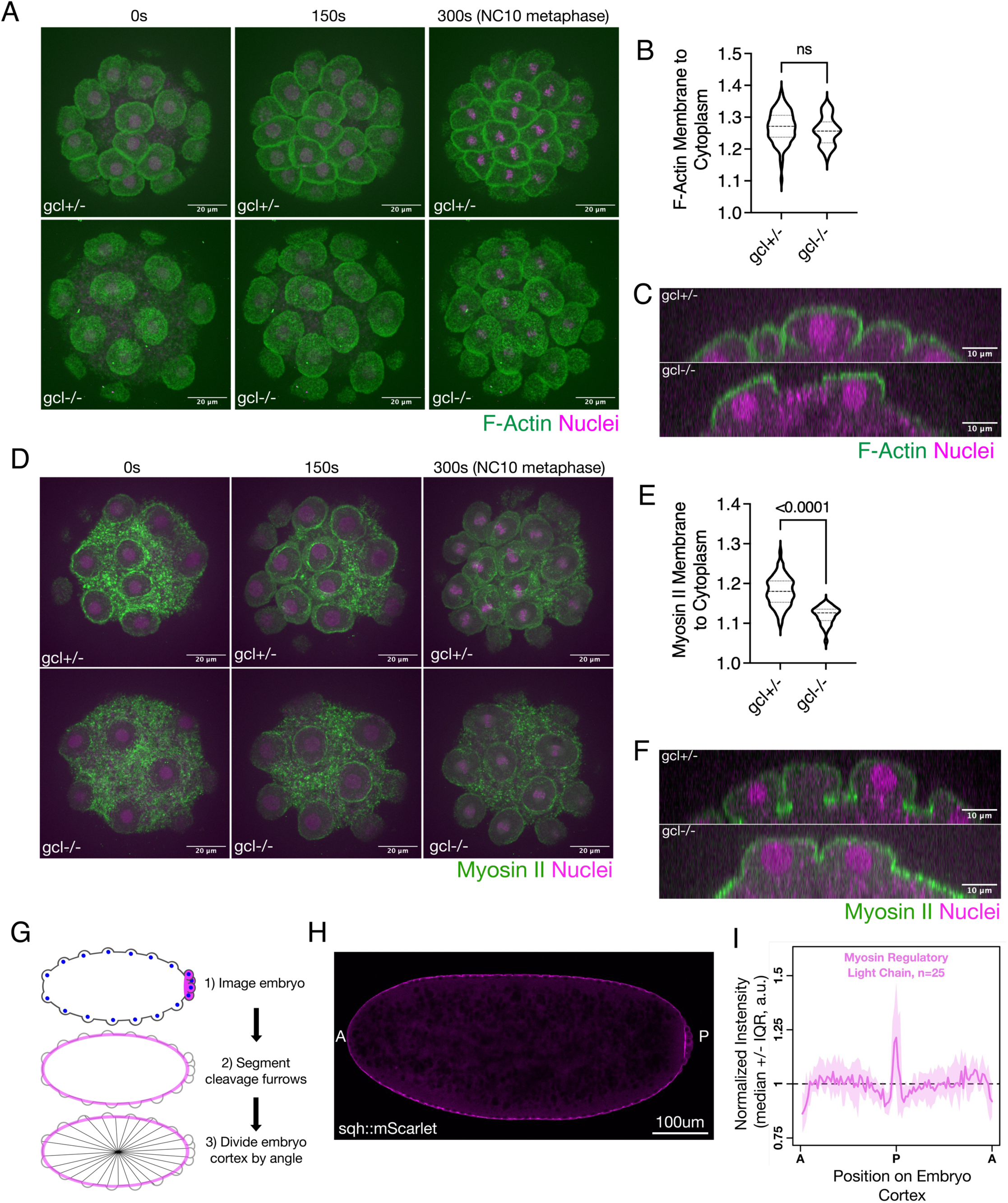
Actin fluorescence is increased in *gcl-/-* pole buds, while myosin II levels are decreased, causing deformation of the pole bud shape. (A) Wild-type and *gcl-/-* embryos expressing the F-actin marker moe[ABD]:GFP (green) and the nuclear marker His2AV:RFP (magenta) were posteriorly mounted and live imaged. Images are maximum intensity projections of a 20 µm section. The 300s time point represents nuclear cycle 10 metaphase. Scale bar = 20 µm. (B) Membrane and cytoplasm were segmented from posterior-mounted embryo movies 150s before nuclear cycle 10 metaphase, and the membrane-to-cytoplasm ratio of F-Actin fluorescence was plotted. (n = 6 embryos for *gcl+/-,* n = 5 embryos for *gcl-/-,* Mann-Whitney test) (C) Orthogonal view of *gcl +/-* and *gcl-/-* embryos expressing the F-actin marker moe[ABD]:GFP (green) and the nuclear marker His2AV:RFP (magenta) 150s before nuclear cycle 10 metaphase. Scale bar = 10 µm. (D) Wild-type and *gcl-/-* embryos expressing a marker for the regulatory light chain of non-muscle myosin II, sqh:3xGFP, (green) and the nuclear marker His2AV:RFP (magenta) were posteriorly mounted and live imaged. Images are maximum intensity projections of a 20 µm section. The 300s time point represents nuclear cycle 10 metaphase. Scale bar = 20 µm. (E) Membrane and cytoplasm were segmented from posterior-mounted embryo movies 150s before nuclear cycle 10 metaphase, and the membrane-to-cytoplasm ratio of Myosin II fluorescence was plotted. (n = 6 embryos for gcl+/-, n = 5 embryos for gcl-/-, Mann-Whitney test) (F) Orthogonal view of *gcl +/-* and *gcl-/-* embryos expressing a marker for the regulatory light chain of non-muscle II, sqh:3xGFP, (green) and the nuclear marker His2AV:RFP (magenta) 150s before nuclear cycle 10 metaphase. Scale bar = 10 µm. (G) Schematic of embryo segmentation (H) Laterally mounted fixed NC11-12 embryo expressing a marker for the regulatory light chain of non-muscle Myosin II, sqh:mScarlet (magenta). Embryo is oriented with its anterior (A) to the left and its posterior (P) to the right. Image shows single plane roughly in the middle of the embryo. Scale bar = 100 µm. (I) Plot of sqh:mScarlett across the circumference of the embryo cortex normalized to the average fluorescence intensity of the cleavage furrows per embryo.

To measure myosin recruitment directly, we live-imaged a GFP-fusion protein of the regulatory light chain of non-muscle myosin II (sqh:GFP), which localizes to regions of active myosin contractility, in control and *gcl-/-* embryos. During nuclear cycle 10, as in somatic syncytial buds, non-muscle myosin II initially formed a diffuse network occupying the cortex between pole buds (Foe et al. 2000; Zhang et al. 2018). About 150 seconds before nuclear cycle 10 metaphase, myosin II organized into ring structures at the base of the pole buds as they began to constrict in control *gcl+/-* embryos while remaining open in *gcl-/-* embryos. The membrane-to-cytoplasm ratio of myosin II fluorescence is higher in *gcl+/-* pole buds compared to *gcl-/-* pole buds, reflecting the direct connection between myosin assembly and pole bud constriction (Fig. 7D-F). The failure of myosin II to localize to the cleavage furrow is consistent with observations in *pten-/- Dictyostelium discoideum* cells, which exhibit ectopic PIP3 production and defective cytokinesis (Janetopoulos et al. 2005).

If myosin II recruitment is inhibited when PIP3 levels are high, we would expect that even in wild-type embryos, myosin II recruitment is decreased at the membrane where Torso RTK activity is high. Indeed, consistent with the spatial patterning of Torso RTK, we found that in fixed embryos, myosin II fluorescence intensity is decreased at the anterior pole and the regions adjacent to the posterior pole, but is increased at the posterior pole where germplasm is recruited, during PGC formation (Fig. 7G-I).

These observations suggest a model in which decreased actomyosin contractility may influence the rounding of pole buds. We measured the height and width of pole buds during cellularization using the membrane marker Katushka2-CAAX in both wild-type and *gcl* mutant embryos. Our findings indicate that the width of pole buds is similar in *gcl-/-* embryos compared to the control, but the height is reduced in *gcl-/-* embryos. Pole buds extend less from the embryo cortex and exhibit a flatter shape in *gcl* mutants (Fig S3A-D). This flat phenotype is consistent with cells lacking PTEN and exhibiting heightened PIP3 levels (Tang et al. 2011), indicating that successful pole bud formation depends on the spatiotemporal regulation of actomyosin contractility through the antagonism between GCL and Torso at the posterior pole.

## Discussion

Here, we describe the role of Torso and its suppressor GCL in establishing a spatial asymmetry in PI3K activation and PIP3 production, which is crucial for proper PGC formation. While Torso’s role in activating the transcription of terminal genes in Drosophila is well established, linking Torso RTK to PI3K activity is unexpected. Our study reveals an early function of Torso RTK activity in the spatial patterning of cortical PIP3 through PI3K activation, which occurs before activation of the Raf/MEK signaling pathway. During nuclear cycle 9-10, nuclei migrate to the cortex, and it is now clear that the membrane they reach already varies in its lipid composition. This also highlights an earlier role of GCL in “preparing” the cortex prior to the arrival of the syncytial nuclei. Once nuclei reach the posterior pole, GCL is recruited to their nuclear envelopes through its NLS, serving as a means to sequester it and prevent further activity (Pae et al. 2017). Since the nuclear localization of GCL is not necessary for its role in protecting PGC formation from Torso RTK (Pae et al. 2017), we propose that GCL’s role in patterning the posterior pole membrane is completed by the time the nuclei arrive.

By targeting Torso for degradation, GCL prepares a membrane domain with low PIP3 levels, thereby facilitating PGC formation in the subset of nuclei that reach the posterior pole. When GCL is not maternally inherited, *gcl-/-* embryos exhibit high PIP3 levels at the posterior pole membrane. Pole buds originating from this PIP3-rich membrane cannot recruit sufficient myosin II for timely bud furrow constriction, preventing PGC formation. It has been suggested that germplasm presence facilitates outward budding produced by polymerization of a branched F-actin network through PIP2 enrichment, although the regulator of this enrichment has not been identified (Kilwein et al. 2025). Asymmetric actomyosin contractility is crucial for establishing polarity in various contexts, such as the early *C. elegans* embryo, where myosin recruitment is inhibited at the posterior pole due to an unknown centrosome-dependent polarity cue. The anterior bias of the actomyosin network correlates with an enrichment of PIP2 cortical structures and depends on the asymmetric distribution of PAR proteins (Scholze et al. 2018). However, further studies revealed that this could be attributed to an asymmetric membrane topology and differential labeling by the membrane probes used (Hirani et al. 2019). Further experiments are needed to show whether a similar asymmetric membrane topology could explain the recently reported posterior enrichment of the PIP2 probe (Kilwein et al. 2025). PIP3 levels are highly tuned to growth factor signaling, while PIP2 levels are not. Since our probes show no difference in PIP2 levels in *gcl+/-* and *gcl-/-* pole buds (Fig. 6B-D), the observed increase in PIP3 synthesis in *gcl-/-* embryos likely results from heightened PI3K activity rather than a simple increase in plasma membrane bunching at the posterior pole. Our study now links the distribution of PIP3 directly to GCL’s action and a restriction of PI3K activity, since it is apparent that even *gcl+/-* pole buds carry more PIP3 than their wild-type counterparts, despite making the same number of PGCs (Fig. 7C). Thus, PGC formation depends on the proper recruitment and spatial distribution of these lipid secondary messengers and their subsequent ability to regulate myosin assembly.

The posterior PIP3-rich membrane domain surrounding germplasm may play a role in limiting the number of PGCs that form, as our findings indicate that inhibiting Ras and PI3K activity causes the emergence of ectopic PGCs, akin to when GCL is overexpressed (Jongens et al. 1994). Ectopic PGCs in embryos overexpressing GCL lack sufficient germplasm factors necessary for survival and specification, resulting in their death before they migrate to the gonad (Slaidina and Lehmann 2017). It may be energetically undesirable for the embryo to generate more germ cells than it can sustain, and the band of PIP3-rich membrane encircling the posterior pole could be one mechanism ensuring that germ cell formation aligns with the available germplasm.

PIP3 levels must be tightly regulated in both space and time to allow cytokinesis to complete, primarily through the opposing behaviors and distinct subcellular localizations of PI3K and PTEN (Janetopoulos et al. 2005). Our findings reveal an additional hierarchy of regulation in the early *Drosophila* embryo that employs the selective degradation of a receptor tyrosine kinase to limit PI3K activity and PIP3 production at the posterior pole. PIP2 and PIP3 levels are known to be important for somatic cellularization in *Drosophila*, and the ability to finely balance myosin assembly and disassembly is essential for furrow growth and constriction later in embryogenesis (Reversi et al. 2014). Outside of the posterior pole, the syncytial buds of the early embryo that give rise to the soma require a balance between actin polymerization and actomyosin contractility to ensure proper swelling (Fernandez-Gonzalez and Harris 2023). Our results demonstrate that a similar equilibrium is necessary for the proper cellularization of posterior pole buds. Failure of myosin recruitment, signaled by PIP3 overproduction, results in flattened pole buds that cannot pinch off from the rest of the embryo. These findings enhance our understanding of this asymmetric cellularization event and the precise coordination of the cytoskeletal machinery required.

It remains unclear whether the creation of Torso-dependent PIP3 domains at the termini of the early embryo has a functional role or is simply a consequence of the initial partitioning of activated Torso during oogenesis, which is necessary to ensure the specification of the terminal regions. *Torso^HH/WK^*embryos do not exhibit any notable defects at nuclear cycle 14, except for the “pole hole,” a cellularization defect at the posterior where germ cells form (Perrimon et al. 1985). Similarly, the overproduction of PIP3 at the posterior pole of *gcl-/-* mutant embryos does not significantly affect nuclear migration or somatic cellularization, but does impair the ability of pole buds to cellularize. Therefore, it can be argued that the interaction between Torso and GCL is essential for establishing a soma-germline border by restricting lipid distribution in the embryonic cortex.

## Acknowledgements

We thank Ben Lin for helping to design germplasm-specific expression constructs and past and present members of the Lehmann lab for helpful suggestions throughout this work. We thank the Bloomington Drosophila Stock Center (NIH P40OD018537) and the Vienna Genome Resource Center, as well as our colleagues Ben Lin, Trudi Schüpbach, Jennifer Zallen, Yohanns Bellaïche, Tom Harris, and Jared Toettcher for fly stocks, and the Drosophila Genome Resource Center (NIH 2P40OD010949) for reagents. We are grateful to the FlyBase consortium for providing data and curators (NHGRI/NIH U41HG000739 and U24HG013300), and the W.M. Keck Microscopy Innovation Center at Whitehead Institute for advice and support. This work was, in part, supported by NIH award (5R01HD110546).

## Author Contributions

Conceptualization: MS, JP, RL; Experimentation and Methodology: MS, JP, AMV, MA; Analysis: MS, JP, AMV; Writing: MJ, RL; Funding Acquisition: RL; Supervision: RL

## Method Details

### D. melanogaster Maintenance

*Drosophila melanogaster* were raised at 25°C on medium containing 1.5% yeast, 3.6% molasses, 3.6% cornmeal, 0.112% Tegosept, 1.12% alcohol and 0.38% propionic acid. Apple juice plates for embryo collection contained 25% apple juice, 2.5% sucrose, 2.25% bactoagar, 0.15% Tegosept.

### Histology

For embryo collection, young parents (less than five days post-eclosure) were transferred to embryo collection cages equipped with yeasted apple juice plates. To optimize embryo collection in the late morning, cages were kept in incubators set to a 10pm-10am light-dark cycle. To collect nuclear cycle 13-14 embryos for PGC counting, flies were allowed to lay eggs for two hours at 25°C. Then the plates were removed, and the embryos were aged for an additional hour before fixation.

Embryos for PGC counting were dechorionated with 50% bleach for two minutes and fixed for 15-20 minutes in a biphasic solution of heptane and 4% PFA-1xPBS on a shaker. The aqueous layer was removed and 10mL of methanol was added for de-vitellenization followed by one minute of robust shaking by hand. Fresh methanol was exchanged three times before storing at -20°C or continuing with immunofluorescence staining protocol. Blocking and staining was carried out in PBS/0.1% bovine serum albumin/0.1% Triton-X.

For myosin II quantification, embryos were dechorionated with 50% bleach for two minutes and fixed for 1 hour in a biphasic solution of heptane and 4% PFA-1xPBS on a shaker, then manually devitellinized as follows: the PFA phase was removed and embryos were rinsed twice briefly with PBS. Using a glass Pasteur pipette, embryos were transferred to plastic Petri dishes and briefly dried to facilitate adhesion to the plastic surface. The embryos were then covered in PBS, and under a dissecting stereomicroscope, manually removed from their vitelline membranes using a fine syringe. PBS/0.3% Triton-X was added to the dish to free the devitellinized embryos from the plastic, and the embryos were transferred to a microfuge tube and washed five times in PBS/0.3% Triton-X to remove all traces of PFA before continuing with the immunofluorescence staining protocol.

### Immunofluorescence

Wash buffer used was PBS/0.1% Triton-X (PBST). All wash steps were performed while embryos rotated periodically on a nutator. Methanol-devitellinized embryos were gradually rehydrated with 3:1, 1:1, and 1:3 Methanol:PBST, then washed three times with PBST before a thirty-minute block with 1% BSA in PBST. Embryos were incubated in primary antibody in blocking buffer overnight at 4°C, using the concentrations listed on the reagents table, and then washed five times with PBST at room temperature. Embryos were blocked again for thirty minutes. They were then incubated in secondary antibody in blocking buffer for four hours at room temperature in the dark, followed by three washes with PBST. Lastly, embryos were equilibrated in Vectashield (Vector Laboratories) overnight at 4°C before mounting on coverslips and imaging.

Immediately after fixation, hand-devitellinized embryos were blocked for 30 minutes in PBS/1% BSA/0.3% Triton X-100, and then counterstained overnight with 1:1000 Phalloidin and 1:2000 DAPI (2mg/mL). Phalloidin (F-actin) and DAPI (DNA) were used for staging embryos. The embryos were washed five times in PBS/0.3% Triton-X, then allowed to equilibrate overnight in Vectashield mounting media. The following day, the embryos were mounted on glass slides and imaged.

Imaging of fixed embryos was done on an inverted Zeiss LSM 780 laser scanning confocal microscope with a Zeiss air Plan-Apochromat 20x/0.8 objective.

### Plasmids

To generate UASp-Ras, UASp-Rgl-nosTCEpgc 3’UTR, UASp-PI3K92E-nosTCEpgc 3’UTR, and UASp-Pi3K21B-nosTCEpgc 3’UTR lines, the coding sequence of the gene of interest was amplified from cDNA clones ordered from DGRC and cloned into a pWallium22 vector containing the PGC-specific 3’UTR (TCEp3), provided by Dr. Benjamin Lin. All constructs were integrated into attp2, attp40, and attp5 by Bestgene. A full list of plasmids is included in the reagents table.

### Timelapse Imaging of PGC Formation

To capture PGC formation, embryos were collected for 1.5 hours. Embryos were dechorionated with 50% bleach for two minutes. Heptane glue was made from dissolving tape adhesive into heptane. To mount for imaging, embryos were glued either on their sides or their posterior poles to 35mm coverslip dishes (MATTEK P35G-1.5-14-C) and submerged with a layer of halocarbon oil 700 (Sigma-Aldrich #H8898). Images were acquired on an inverted Nikon Ti2 with a Yokogawa CSU W1 spinning disk scanhead and Kinetix camera, using a Nikon Plan Apo Lambda D DIC N2 60x/1.42 oil objective controlled by Nikon Elements. For posterior-mounted embryo time-lapse movies, stacks of 20 µm thickness with 1 µm step size were acquired every 30 seconds. All stacks begin at the coverslip in order to capture the entire membrane of the pole buds.

Images for pole bud height and width measurements (Fig. S3) were acquired on an upright LSM 980 laser scanning confocal microscope with a Zeiss air Plan-Apochromat 20x/0.8 objective. To mount for imaging on an upright scope, embryos were glued on their sides to a coverslip and placed on a gas-permeable membrane (SARSTEDT #94.6170.102) that was covered with halocarbon oil 27 (Sigma-Aldrich #H8733).

### Optogenetic Activation of Sos

Transgenic flies carrying the OptoSos construct were kindly provided by the Toettcher laboratory (Johnson et al., 2017). Embryos were collected on apple juice plates for 30 minutes, dechorionated in 50% bleach for 2 minutes, thoroughly rinsed in water, and then mounted on coverslips using heptane glue. Embryos were imaged and activated using an inverted Zeiss LSM 780 laser scanning confocal microscope with a Zeiss air Plan-Apochromat 20x/0.8 objective. Light (488nm) was applied at 40% power for 10 iterations every 30 seconds starting 10 minutes after collection for 100 minutes.

### PIP3/PIP2/F-actin/Myosin II Pole Bud Measurements

Fluorescence intensity analysis of PIP3, PIP2, F-actin, and Myosin II in time-lapse images of posteriorly mounted live embryos (Fig. 6C, 6F, 7C, 7D, 8B, 8E) was performed using custom Python scripts. Prior to segmentation, the nuclear marker channel of the two-channel z-stack fluorescence images was denoised using a Noise2Void (N2V) (Krull et al., 2019) model to enhance signal-to-noise ratio, trained from images from timepoints not included in the final analysis. Subsequently, pole buds were segmented in 3D using a Cellpose-SAM model (Pachitariu et al., 2025), and nuclei were segmented using a Cellpose cyto3 model (Stringer and Pachitariu, 2025). Both models were trained following a human-in-the-loop approach (Pachitariu and Stringer, 2022) using random xy slices from the datasets and 3D segmentations generated using the stitching mode in Cellpose.

The resulting 3D masks for pole buds and nuclei were then processed to define three distinct cellular compartments: nucleus, cytoplasm, and cell membrane. The nuclear compartment was directly defined by the nucleus mask. To define the pole bud periphery for membrane analysis, whole pole bud masks underwent subtraction of a 3D erosion operation from a dilation operation, both using a ball structuring element. This peripheral mask was further refined by thresholding, retaining only pixels with an intensity in the PIP3/PIP2/F-actin/Myosin II channel greater than 105% of the mean cytoplasmic intensity. The mean cytoplasmic intensity for this refinement step was measured in a region defined by dilating the nucleus mask by 2 pixels (0.2 µm). The cytoplasm mask was generated by eroding the whole cell mask and subsequently subtracting the nuclear mask.

For each segmented pole bud, mean PIP3/PIP2/F-actin/Myosin II intensities within the nucleus, cytoplasm, and cell membrane masks were calculated from raw images without additional pre-processing. Intensity ratios between membrane and cytoplasm were determined.

Figures and plots were generated using ImageJ and GraphPad Prism. The Mann-Whitney U test was used on GraphPad to compare data sets. Every time point from the timelapse was processed using the method described above.

### PGC Counting

Fixed embryos were stained with anti-Vasa antibody and DAPI to visualize pole cells and DNA. Only embryos that were at nuclear cycle 13-14, prior to somatic cellularization, were counted. Multiple z-planes with a 2.5um step size encompassing the area of PGC formation were imaged on an inverted Zeiss LSM 780 laser scanning confocal microscope with Zeiss air Plan-Apochromat 20x/0.8 objective. Vasa-positive PGCs in each embryo were counted manually using the ImageJ Cell Counter plugin (NIH; http://rsb.info.nih.gov/ij/) (Schindelin et al., 2012). The Mann-Whitney U test was used on GraphPad to compare data sets.

### Myosin II Quantification in Fixed Embryos

The following imaging and analysis protocol was performed on hand-devitilinized embryos expressing sqh-sqh:mScarlet. Single z-slices containing the middle Z-plane of each embryo were acquired on a Zeiss 780 point scanning confocal microscope with 2x averaging, 16bit depth and 2048x2048 image size using the 561 nm laser line with 0.6x zoom. Zeiss air Plan-Apochromat 20x/0.8 objective was used. Images were then manually rotated to align the A-P axis horizontally.

To quantify myosin signal profiles across the embryo circumference, masks of the myosin signal were segmented and intensity was quantified using a custom ImageJ macro as follows. First, cleavage furrows were emphasized using the find edges function; then, a pixel-maximum filter with a three-pixel radius was applied. Next, a binary threshold was manually applied to create a mask that encompassed the embryonic cleavage furrows. Gaps in the detected contours of the embryo circumference were manually filled to produce a single, continuous mask representing the embryo circumference. The ROI corresponding to the mask was then transferred to the original unaltered image for quantification. The ROI was radially divided into 100 equal-angle segments and the mean myosin signal intensity for each segment was quantified. These data were then processed in R to normalize each segment’s average myosin signal intensity to the average myosin signal across the entire ROI for each embryo, which allowed for combining myosin intensity profiles from multiple embryos in a single plot. Plots were generated using the base plotting system in R.

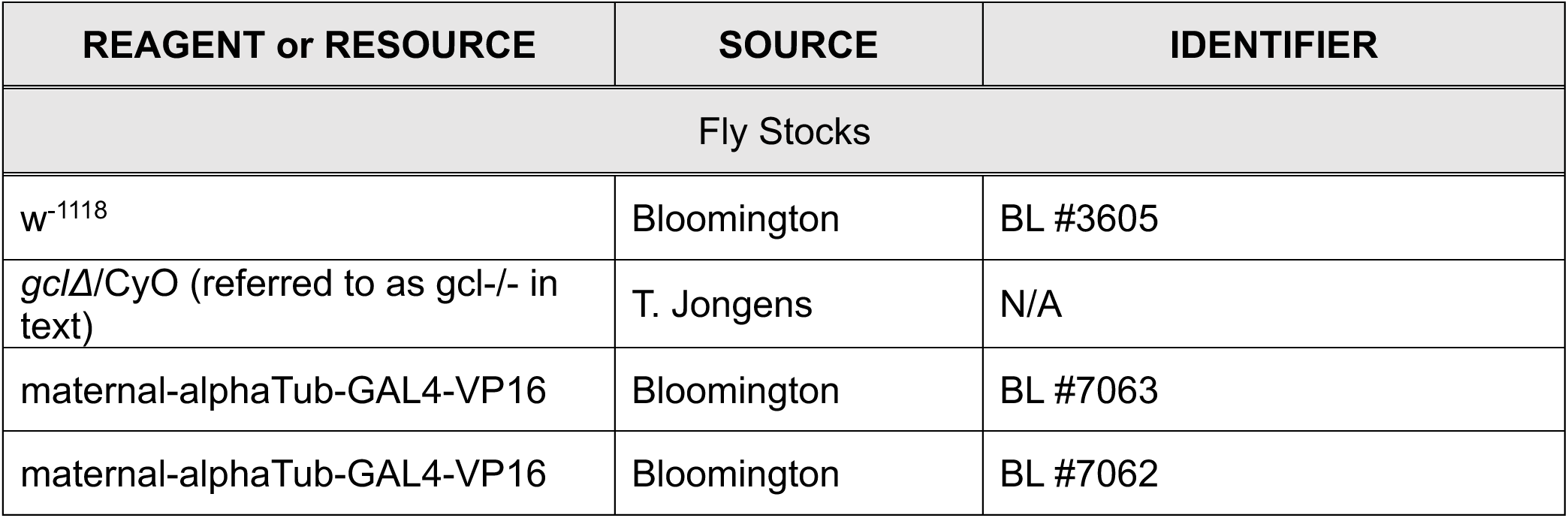
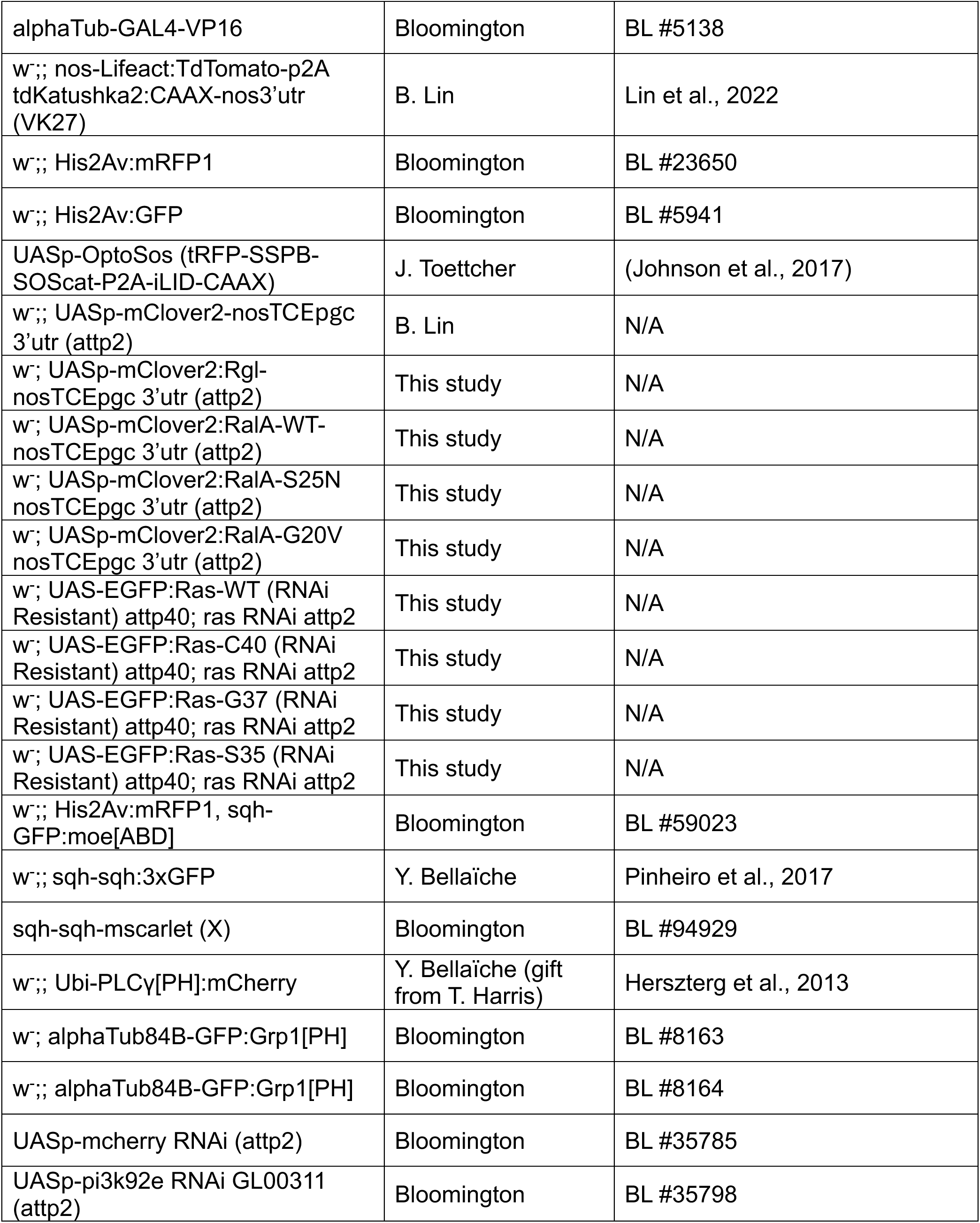
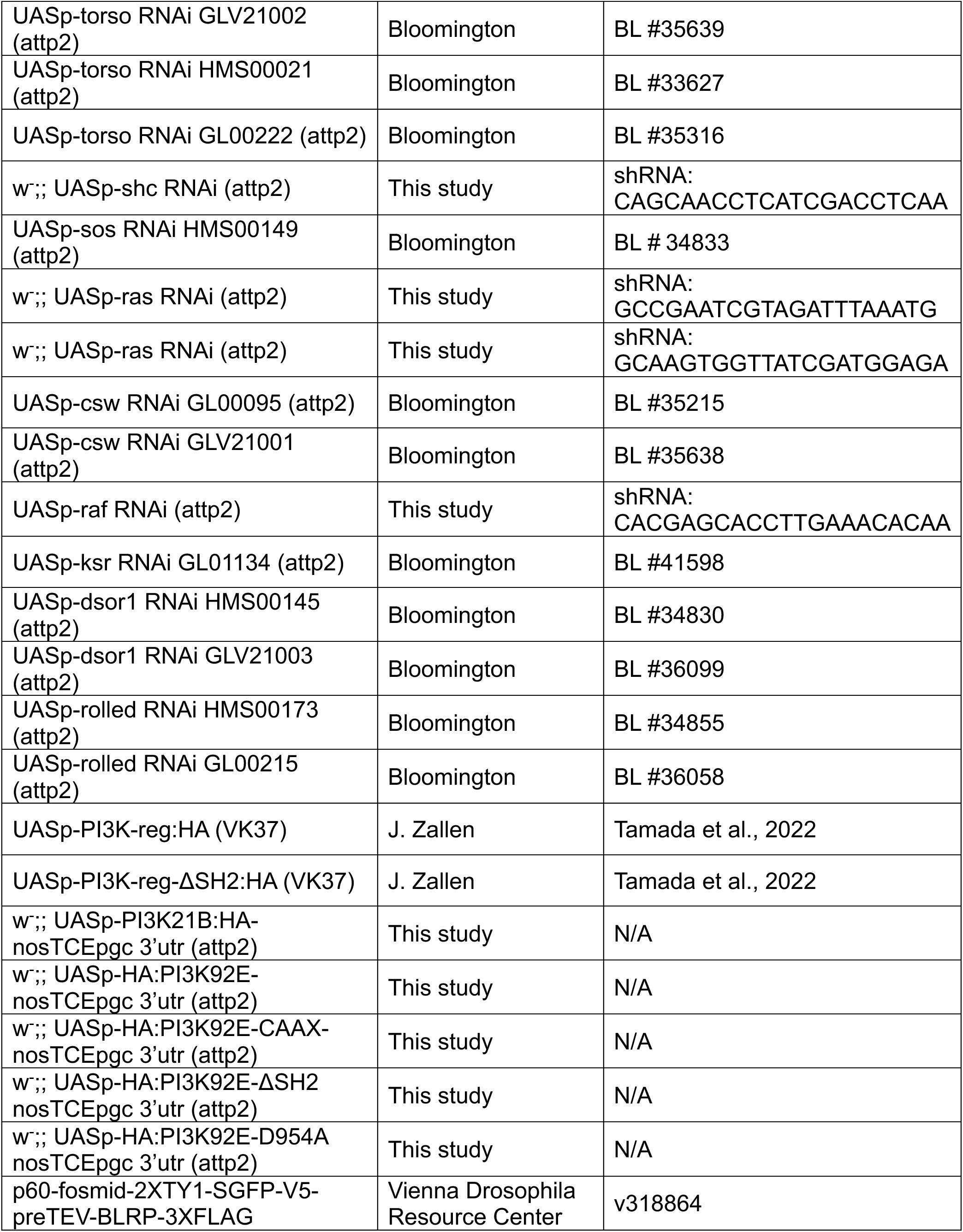
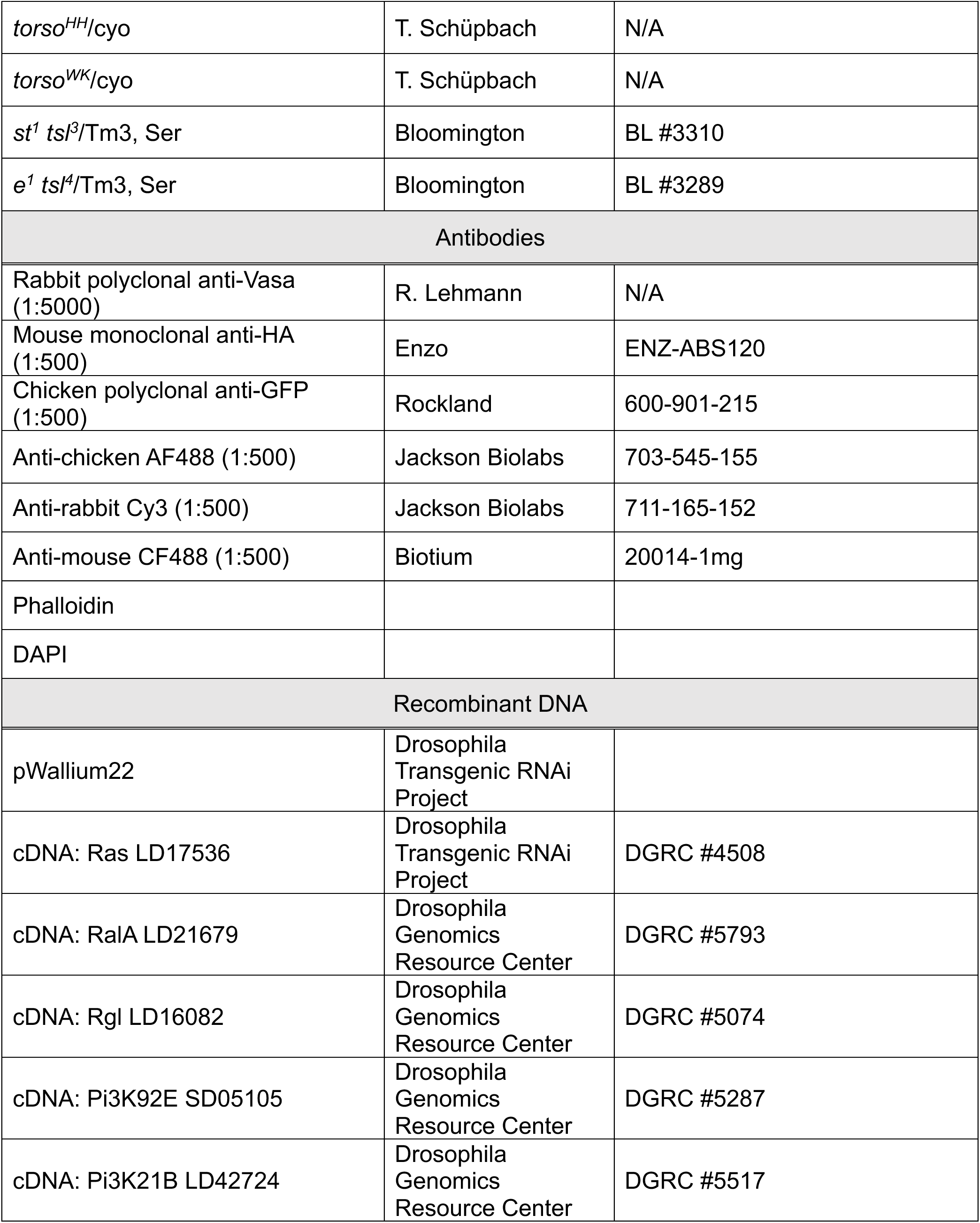
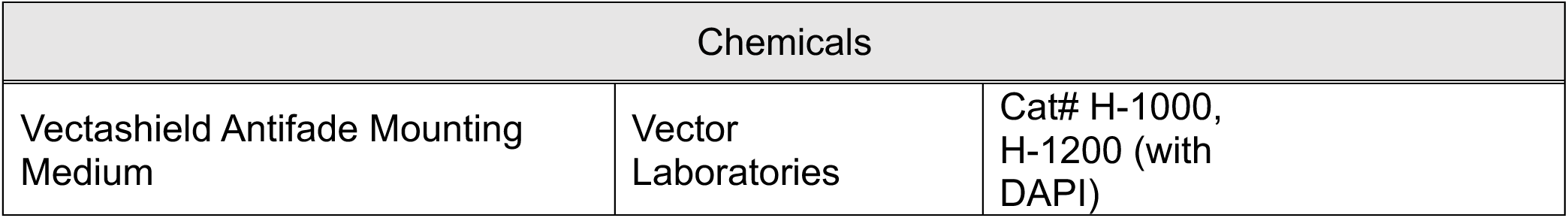

