## Supplemental Figures for "GCL pruning of PIP3 establishes the soma-germline boundary"

#### Supplementary Figures

### Supplementary Figure 1

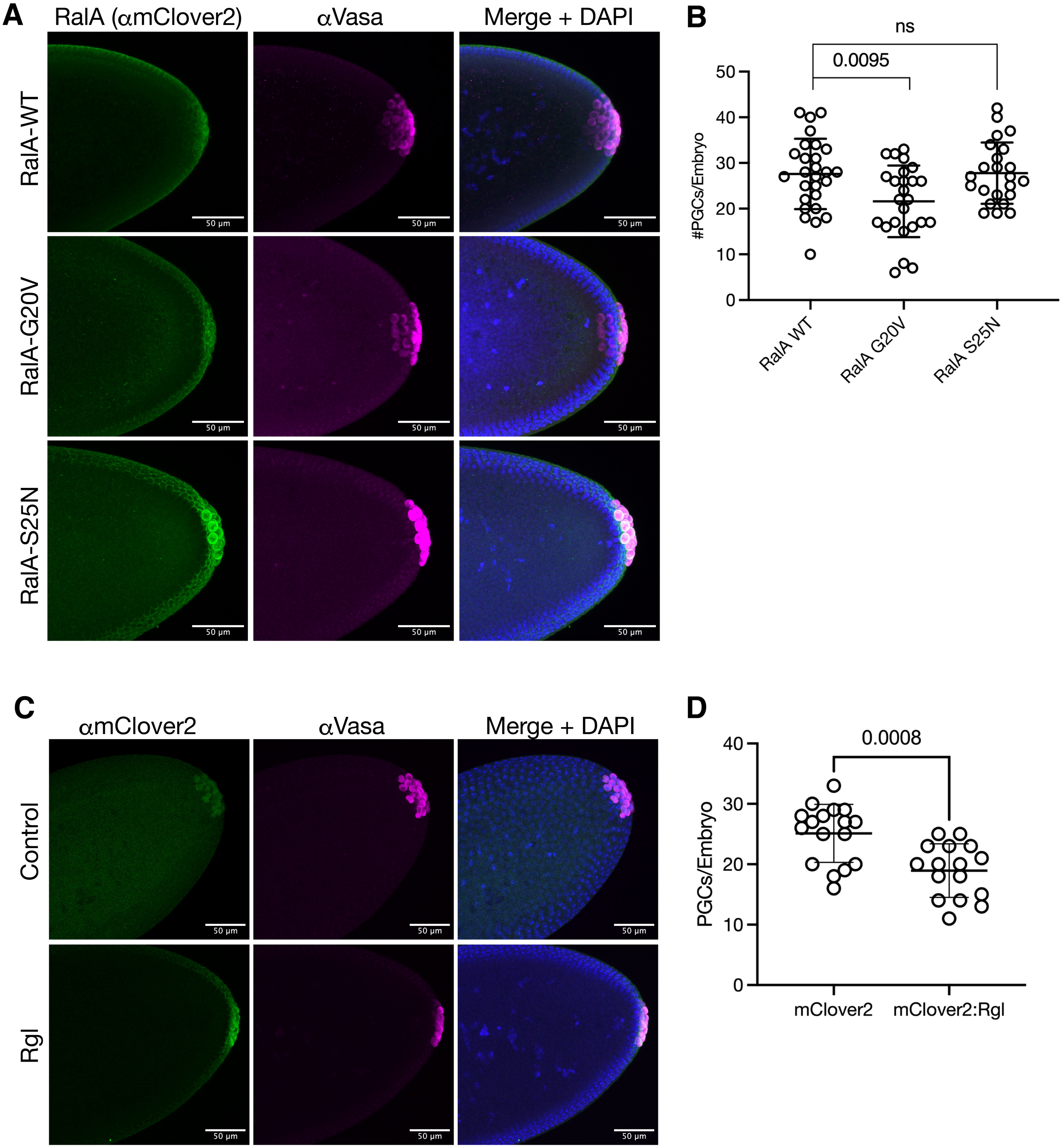

**Supplementary Figure 1: Increasing RalA activity mildly antagonizes PGC formation**

(A) Embryos overexpressing germline-targeted, mClover2-tagged RalA-WT, RalA-G20V (constitutively active), and RalA-S25N (dominant negative) were immunostained with anti-mClover2 and anti-Vasa to confirm expression of the construct and stain PGCs for counting. Expression of the UASp constructs was driven by two copies of maternal-tubulin-GAL4-VP16 for optimal expression during late oogenesis. Images depict maximum intensity projections spanning area of PGC formation. Scale bar = 50 μm.

(B) (B) PGCs in nuclear cycle 13-14 embryos from mothers of the indicated genotype were counted and plotted from (A). Overexpression of RalA-WT was used as a control. Bars represent the mean ± standard deviation. (n > 20, Mann-Whitney test)

(C) Embryos overexpressing germline-targeted, mClover2-tagged Rgl were immunostained with anti-mClover2 and anti-VASA to confirm expression of the construct and stain PGCs for counting. Expression of the UASp construct was driven by two copies of maternal-tubulin-GAL4-VP16 for optimal expression during late oogenesis. Images depict maximum intensity projections spanning the area of PGC formation. Scale bar = 50 μm.

(D) Number of PGCs in nuclear cycle 13-14 embryos from mothers of indicated genotype was counted and plotted from (B). Overexpression of germline-targeted mClover2 was used as a control. Bars represent the mean ± standard deviation. (n > 15, Mann-Whitney test)

### Supplementary Figure 2

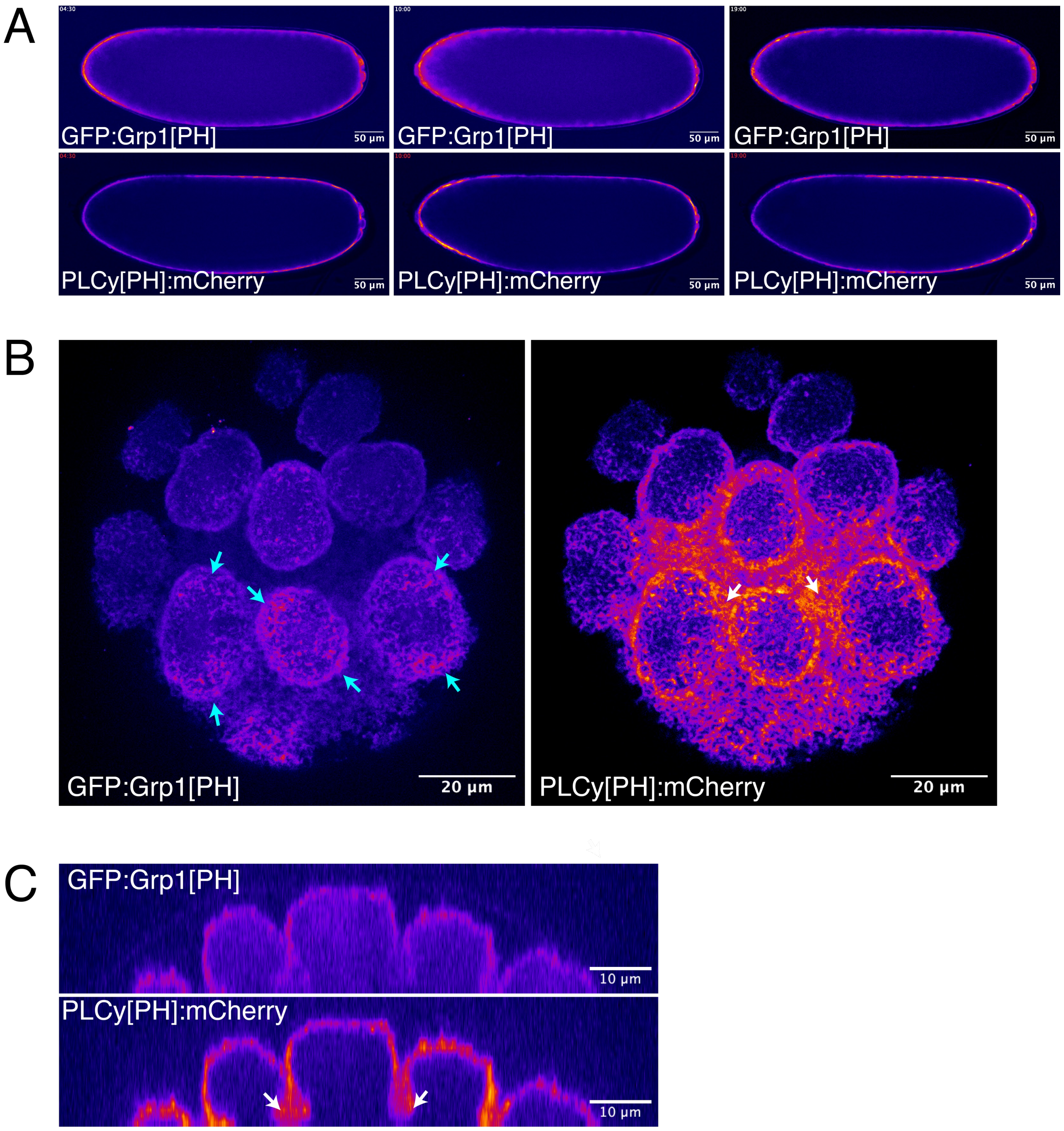

**Supplementary Figure 2: PIP2 and PIP3 occupy separate membrane compartments of the embryo and the posterior pole**.

(A) Embryos expressing the PIP3 biosensor GFP:Grp1[PH] and the PIP2 biosensor PLCγ[PH]:mCherry were live imaged, mounted laterally. Images show a single plane approximately midway through the embryo at three time points during nuclear cycle 10-11, prior to PGC formation. Scale bar = 50 µm.

(B) Embryos expressing GFP:Grp1[PH] and PLCγ[PH]:mCherry were live imaged, mounted on its posterior pole. Embryo is nuclear cycle 10-11, prior to PGC formation. Timing is approximately five minutes before mitosis. Enrichment of PIP3 biosensor seen at poles of buds (cyan arrows). PIP2 is enriched in the interbud regions (white arrows). The image shows the maximum intensity projection of a 20 µm section. Scale bar = 20 µm.

(C) Orthogonal view of embryo from (B) to visualize pole bud furrows. PIP2 but not PIP3 is enriched at the bud furrow (white arrows). Scale bar = 20 µm.

Supplementary Figure 3

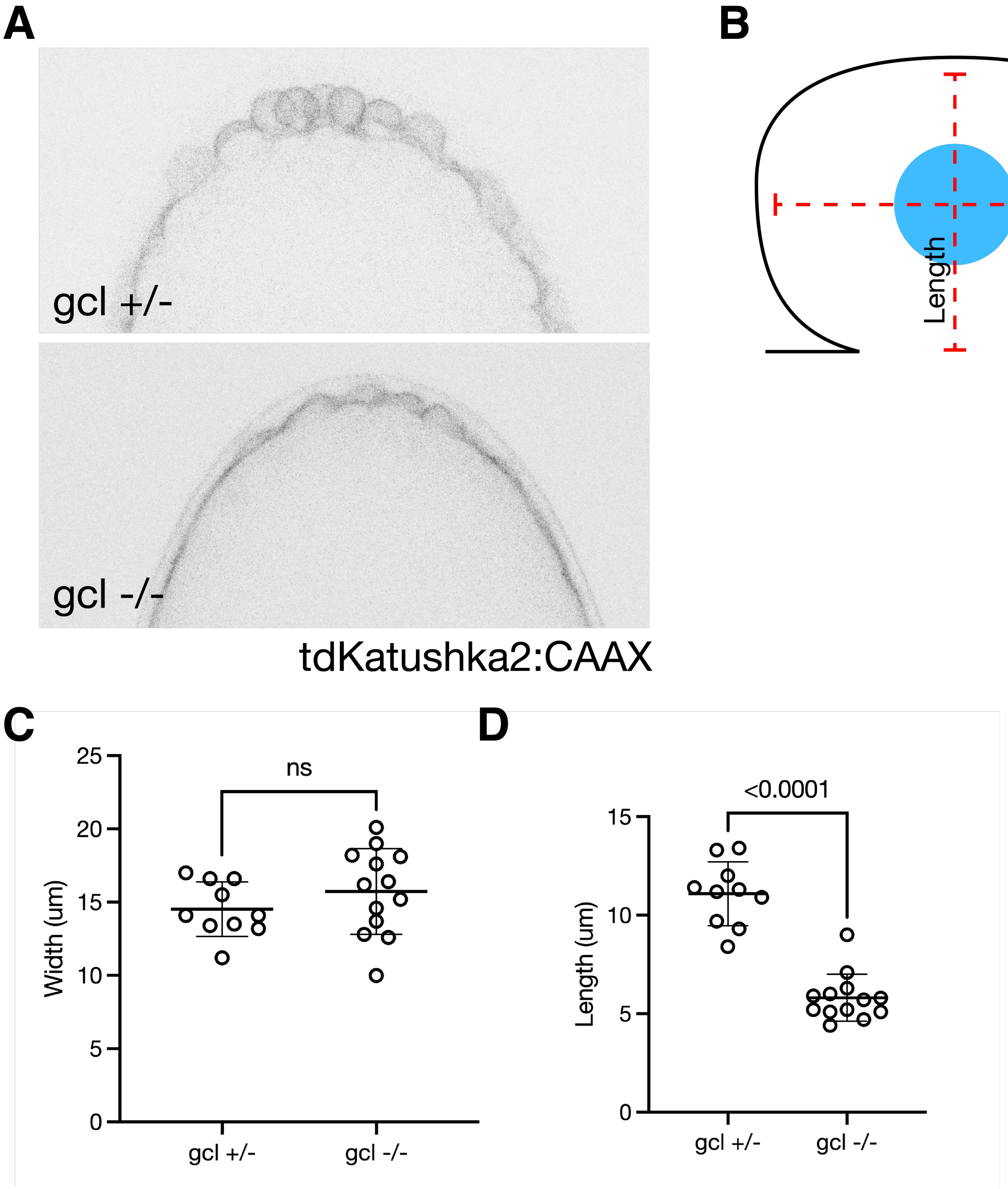

**Supplementary Figure 3: Pole bud membrane shape in *gcl-/-* embryos is flattened due to phosphoinositide imbalance**

(A) Nuclear cycle 10 *gcl*+/- and *gcl*-/- embryos expressing the membrane marker tdKatushka2:CAAX were live imaged on their side. These pole buds are tightly clustered together but have not cellularized. Scale bar = 20 µm.

(B) Diagram of pole bud length and width measurements.

(C) The width of pole buds was measured and plotted. (n = 10 buds for *gcl +/-,* n = 13 buds for *gcl-/-,* Mann-Whitney test, Bars represent the mean ± standard deviation. )

(D) The length of pole buds was measured and plotted. (n = 10 buds for *gcl +/-,* n = 13 buds for *gcl-/-,* Mann-Whitney test, Bars represent the mean ± standard deviation. )

Supplementary Figure 4

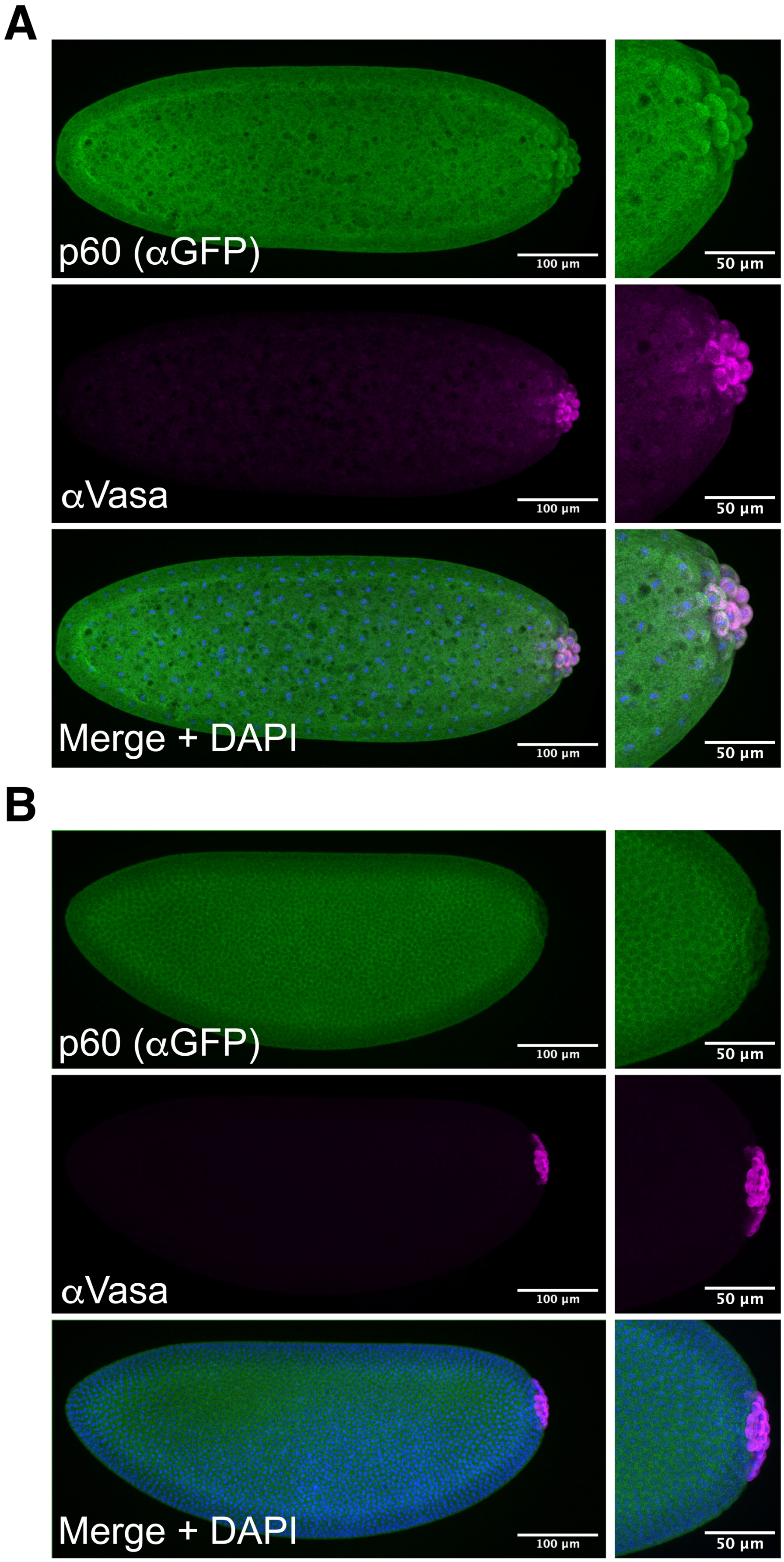

**Supplementary 4: P60, the regulatory subunit of PI3K in *Drosophila*, is uniformly distributed across the embryo before PGC formation, but by nuclear cycle 14 localized to furrows and is excluded from PGCs.**

(A) Nuclear cycle 9-10 embryos shortly preceding pole cell formation expressing p60 fosmid were immunostained for anti-GFP and anti-Vasa. Inset shows a close-up of posterior pole. Images depict maximum intensity projections spanning area of PGC formation. Scale bar = 100 µm for whole embryo, 50 µm for inset

(B) Nuclear cycle 13-14 embryos after pole cell formation expressing p60 fosmid were immunostained for anti-GFP and anti-Vasa. Inset shows a close-up of posterior pole. Images depict maximum intensity projections spanning area of PGC formation. Scale bar = 100 µm for whole embryo, 50 µm for inset.

Supplementary Movie 1

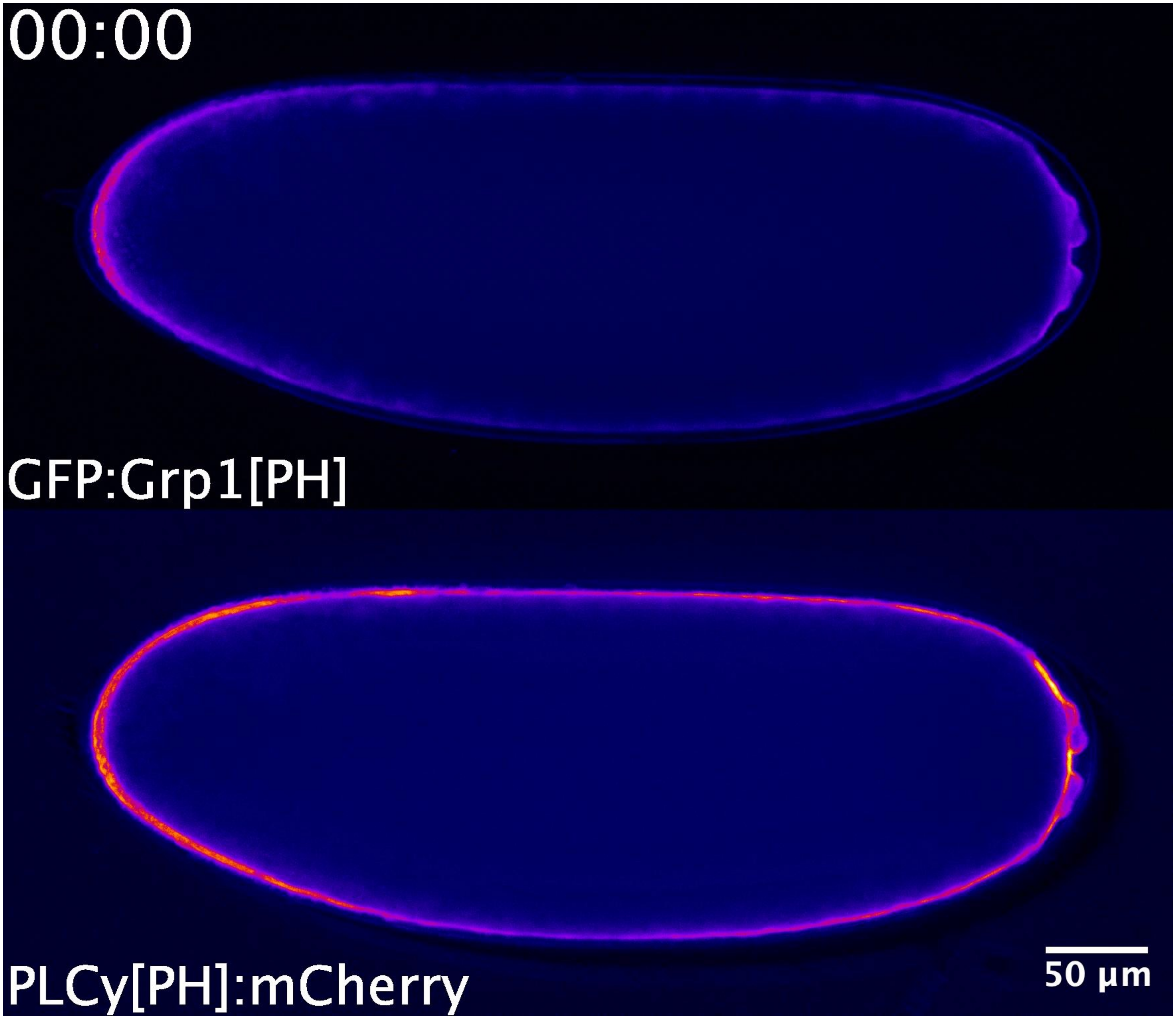

**Movie 1: Embryo expressing the PIP3 biosensor GFP:Grp1[PH] and the PIP2 biosensor** PLCγ[PH]:mCherry was live imaged every thirty seconds, mounted laterally. The embryo is in nuclear cycle 10-11, prior to PGC formation. The image shows a single plane approximately midway through the embryo. Scale bar = 50 μm
